# Chirality-Driven Photonic PCR Enables Label-Free Genetic Amplification

**DOI:** 10.64898/2026.10.19.760185

**Authors:** Haeun Kang, Jeongmin Park, Hyung Joo Lee, Jeong Hyun Han, Youngbi Kim, Dong-Il Won, Donghyun Kim, Mijin Kwon, Dabin Kim, Pil J. Yoo, Jeong Woo Han, Dongseok Suh, Ki Tae Nam, Chul Hoon Kim, Luke P. Lee, Dong Ha Kim

## Abstract

Photonic polymerase chain reaction (PCR) offers a rapid alternative to conventional thermal cycling for genetic amplification, but current platforms are limited by low photothermal conversion efficiencies and a dependence on bulky fluorescent monitoring systems^1–6^. Here we report a chirality-driven photonic amplification paradigm that achieves both accelerated nucleic acid replication and direct, label-free detection. By leveraging chiral plasmonic nanoparticles, we exploit their elevated absorption cross-sections to achieve exceptional light-to-heat conversion and rapid thermal relaxation, significantly accelerating heating and cooling rates during cycling. We show that localized chiral near-fields generated by these chiral plasmonic nanoparticles optimize DNA polymerase conformational dynamics, lowering the activation energy barrier for synthesis and boosting amplification efficiency. Furthermore, by tracking the reaction with circular dichroism spectroscopy, we directly monitor the intrinsic chiroptical signature of amplifying DNA, completely eliminating the need for external fluorescent dyes or labeled probes. We validate this system using SARS-CoV-2 E-gene plasmid DNA templates, demonstrating rapid, label-free amplification and single-copy detection within clinically relevant timelines. By bridging asymmetric photothermal dynamics with chiral-field-induced enzymatic activation, this work translates chiroplasmonic materials from fundamental physics into high-performance, point-of-care molecular diagnostics.

## Introduction

The rapid and accurate replication of nucleic acids underlies modern molecular diagnostics, biosensing, and therapeutic monitoring. For decades, PCR has served as the gold standard for genetic amplification^7–12^, yet conventional instrumentation remains fundamentally limited by the thermal inertia of metal blocks and Peltier elements, which require substantial time to complete sequential denaturation and annealing cycles^13,14^. Photonic PCR has emerged as an alternative that circumvents these macroscopic heat-transfer delays by exploiting the localized light-to-heat conversion of plasmonic nanoparticles to drive rapid thermal cycling^15–22^. Despite this conceptual advance, most current photonic PCR systems still rely on achiral light-matter interactions, where nanoparticle geometry primarily enhances broadband optical absorption or heat generation^4,15,23–26^. Consequently, the distinct roles of optical and structural chirality in governing photothermal amplification remain critically understudied. More critically, these platforms continue to rely on conventional fluorescence-based detection, using intercalating dyes or fluorophore-labelled probes^27–31^. This reliance on fluorescence introduces well-known drawbacks, including photobleaching, background signal interference, and the need for complex optical filtering, all of which constrain the miniaturization and deployment of point-of-care diagnostic systems^32–34^.

Chiral plasmonic nanostructures provide a unique opportunity to overcome these limitations. Because of their broken mirror symmetry, chiral plasmonic nanoparticles exhibit extraordinary optical asymmetry, yielding distinct extinction cross-sections and near-field distributions for left-handed circularly polarized light (LCP) versus right-handed circularly polarized light (RCP)—an optical property fundamentally inaccessible to achiral counterparts^34–45^. Under chirality-matched circular polarization, this asymmetry concentrates absorbed optical energy into highly localized near-field hotspots, offering a structural design parameter for enhancing light-to-heat conversion and its subsequent dissipation during photothermal PCR cycling. While various chiral nanostructures, such as gold helicoids, have been successfully synthesized and investigated for their fundamental physical properties, their translation into these specific functional opportunities remains largely unexplored: most studies of chiral plasmonics to date have focused on enantioselective sensing^39^ or catalysis^43^, rather than on genetic amplification or molecular diagnostics. Consequently, the chiroplasmonics community has frequently faced critical questions regarding the practical utility and immediate real-world benefits of these sophisticated materials. Here, we present a paradigm-shifting aspect of chiral nanomaterials that directly answers this call, demonstrating that engineered chiroplasmonic platforms can radically redefine the boundaries of clinical diagnostics by driving ultra-rapid, label-free genetic amplification.

In this work, we present a chirality-driven photonic amplification strategy that overcomes the limitations of current diagnostic platforms, enabling rapid, label-free detection of pathogenic genomic markers on highly compressed timelines. Moving far beyond a simple iterative application of existing nanoparticles, this study probes and uncovers fundamental, previously invisible physical and biochemical phenomena. First, we exploit asymmetric excited-state relaxation and ultrafast heat-dissipation pathways in chiral gold nanoparticles (AuNPs) under circularly polarized light (CPL) irradiation to overcome the photothermal constraints of conventional achiral systems. This asymmetric photothermal efficiency enables rapid, precise temperature oscillations within compressed cycling windows. Second, we unveil an extraordinary biophysical mechanism wherein the intensely localized chiral near-fields do not merely act as localized heat sources but actively interact with the complex macromolecular machinery of amplification, creating an energetically favorable environment that accelerates the critical conformational transitions of *Taq* DNA polymerase during nucleotide integration. Finally, by eliminating external fluorescent dyes, this framework captures the direct structural transition of replicating DNA within CD measurement setup, thereby directly reading out the native chiroptical signals from the DNA backbone itself^46^. By providing a label-free, ultra-rapid molecular readout, this system offers a versatile blueprint for next-generation point-of-care testing, promising to substantially accelerate and streamline decentralized clinical diagnostics.

## Results

### Chiral plasmonic design for rapid and label-free photonic PCR

We designed a photonic PCR platform in which CPL is coupled to chiral plasmonic AuNPs to induce rapid thermal cycling within a confined reaction volume (**Fig. 1a and Extended Data Fig. 1,2**). The reaction mixture contained chiral AuNPs, Taq DNA polymerase, deoxynucleotide triphosphates (dNTPs) and DNA templates, allowing optical energy to be converted locally into heat while preserving the biochemical components required for polymerase-mediated amplification. Within this system, the chiral AuNPs function as photothermal transducers. Upon absorbing incident CPL, their asymmetric plasmonic geometry generates localized photothermal fields, which induce the periodic temperature changes required for denaturation, annealing and extension without the need for a conventional physical heating block. Because the chiral plasmonic structures interact asymmetrically with CPL, we reasoned that structural chirality could serve as an additional optical design parameter for accelerating photonic PCR beyond what is achievable with achiral photothermal heating.

**Figure 1.**
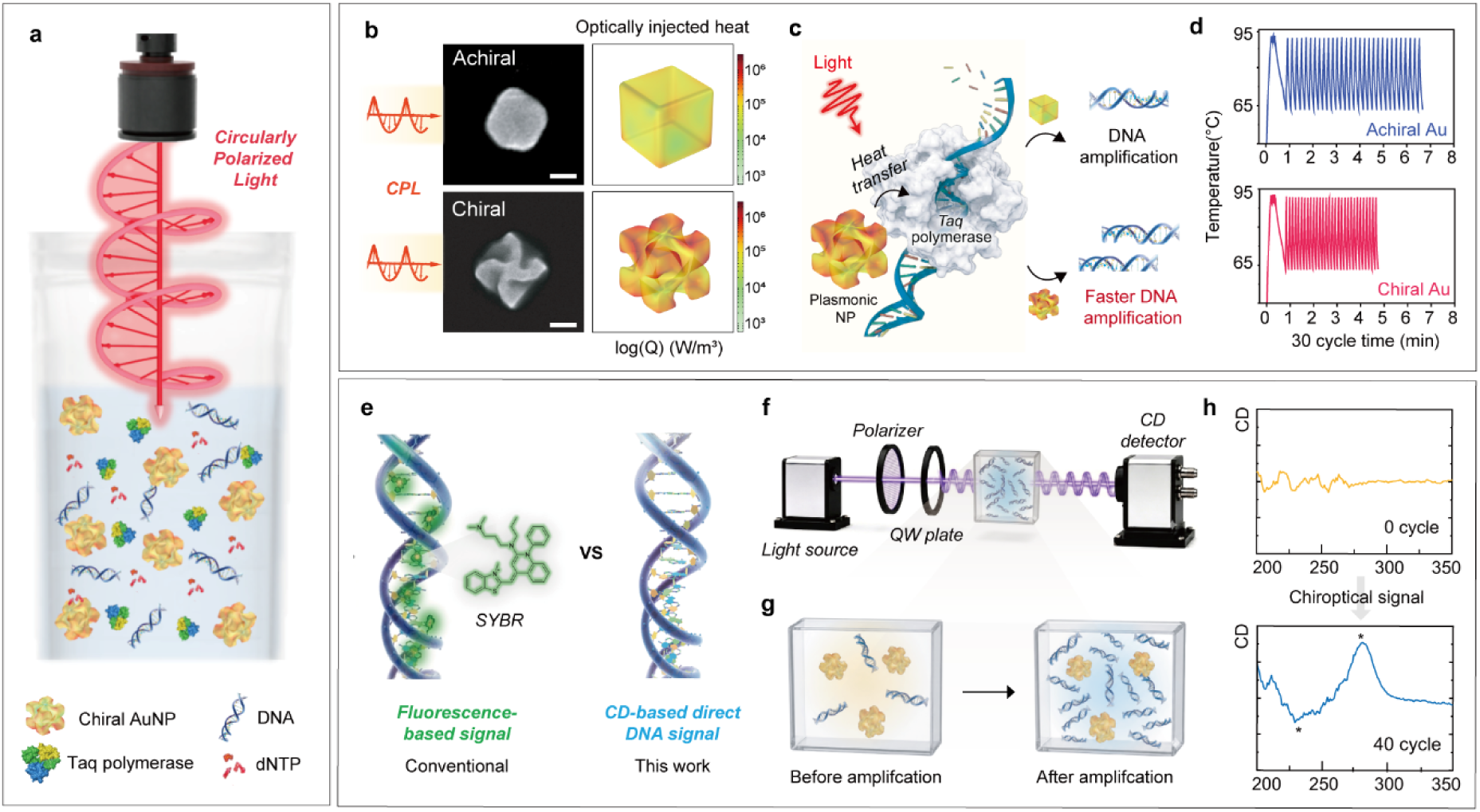
Label-free photonic PCR platform based on chiral plasmonic NPs and CPL. (a) Schematic illustration of the photonic PCR system, in which CPL irradiates a reaction solution containing chiral AuNPs, *Taq* polymerase, dNTPs, and DNA templates. Chiral AuNPs convert incident light into localized heat, enabling thermocycling for DNA amplification. (b) Representative electron microscopy images of achiral and chiral AuNPs and simulated maps of optically injected heat, showing stronger localized heat generation from the chiral nanostructure under CPL irradiation. (c) Schematic illustration comparing heat transfer from achiral and chiral AuNPs during photonic PCR. Chiral AuNPs convert light to heat more efficiently than achiral counterparts, promoting *Taq* polymerase-mediated DNA amplification. (d) Representative temperature profiles recorded during 30-cycle photonic PCR using achiral and chiral AuNPs, showing faster thermal oscillation in the chiral Au system. (e) Comparison between conventional fluorescence-based PCR detection using SYBR dye and the label-free detection strategy used in this work, which directly reads the DNA amplification signal through chiroptical changes. (f) Schematic of the CD measurement setup used to monitor the amplification process. (g) Illustration of the PCR sample before and after amplification, showing increased amplified DNA products. (h) CD spectra recorded before amplification (0 cycle) and after amplification (40 cycles). The amplified sample exhibits a pronounced chiroptical response, demonstrating label-free readout of DNA amplification through the intrinsic CD signal of amplified DNA.

To examine the physical basis of this design, we compared chiral and achiral AuNPs of matched composition under identical CPL irradiation. Representative scanning electron microscopy (SEM) images confirmed the distinct morphologies of the achiral and chiral nanostructures, while simulated maps of optically injected heat showed a more spatially localized and intensified heat-generation profile for the chiral AuNPs relative to the achiral particles under matched optical input (**Fig. 1b**). This enhanced photothermal localization is expected to provide a direct advantage for enzymatic amplification by rapidly modulating the thermal microenvironment surrounding the reaction components (**Fig. 1c**). Consistent with this design principle, the chiral AuNP system generated repeated temperature oscillations over 30 PCR cycles within a shorter cycle window than the achiral AuNP system under matched irradiation conditions (**Fig. 1d**). This comparison demonstrates the central operational advantage of the chiral photonic PCR platform: structural chirality enables improved thermal cycling kinetics beyond those accessible with conventional achiral photothermal heating.

We next integrated a label-free optical readout into the same platform by directly monitoring the chiroptical response of the amplified DNA itself, rather than relying on an external reporter. Conventional PCR detection commonly relies on fluorescent intercalating dyes such as SYBR, which report DNA accumulation indirectly through dye–DNA binding. Because DNA is itself a chiral biopolymer, its amplification can instead be monitored through its intrinsic circular dichroism (CD) signal, without requiring any exogenous label (**Fig. 1e**). We therefore designed a CD-based detection strategy that directly reads out the amplified DNA product without introducing external fluorescent dyes or labelled probes (**Fig. 1f**). Before amplification, the reaction solution contains only a small number of DNA copies, resulting in a weak chiroptical response. After amplification, the increased concentration of DNA products produces a stronger CD signal arising from the helical backbone and base-stacking structure of DNA (**Fig. 1g,h**). These results demonstrate that the same chirality-based platform supports both rapid photonic amplification and direct label-free optical detection of amplified DNA products.

### CPL-induced plasmonic energy localization and heat dissipation

To elucidate the physical origin of the enhanced photothermal conversion observed in the chiral photonic PCR system, we first performed a finite element method (FEM)-based wave optics simulations using commercial Maxwell equation solver (COMSOL Multiphysics), for the AuNPs under CPL illumination to evaluate the optical response of the excited nanostructures. For the chiral AuNP, the differential electric-field map revealed a spatially asymmetric and highly localized field distribution along the helicoidal geometry (**Fig. 2a**). By contrast, the achiral AuNP exhibited a weaker and more spatially symmetric differential response under the same excitation condition (**Fig. 2b**). These simulations suggest that the chiral plasmonic architecture amplifies the near-field imbalance between opposite circular polarizations, creating localized optical environments that are not accessible to achiral particles. Calculated absorption cross-sections further showed that the chiral AuNPs exhibited stronger absorption under both CPL conditions than the achiral AuNPs at 532 nm wavelength (**Fig. 2c**). Experimental absorption spectra obtained using randomly polarized light indeed revealed comparable plasmonic bands in the visible region (**Fig. 2d**). Although the CPL-dependent difference in absorption for the chiral AuNPs was weakly resolved in the simulations, the enhanced photo-selection under chirality-matched CPL was observed in the time-resolved measurements described below.

**Figure 2.**
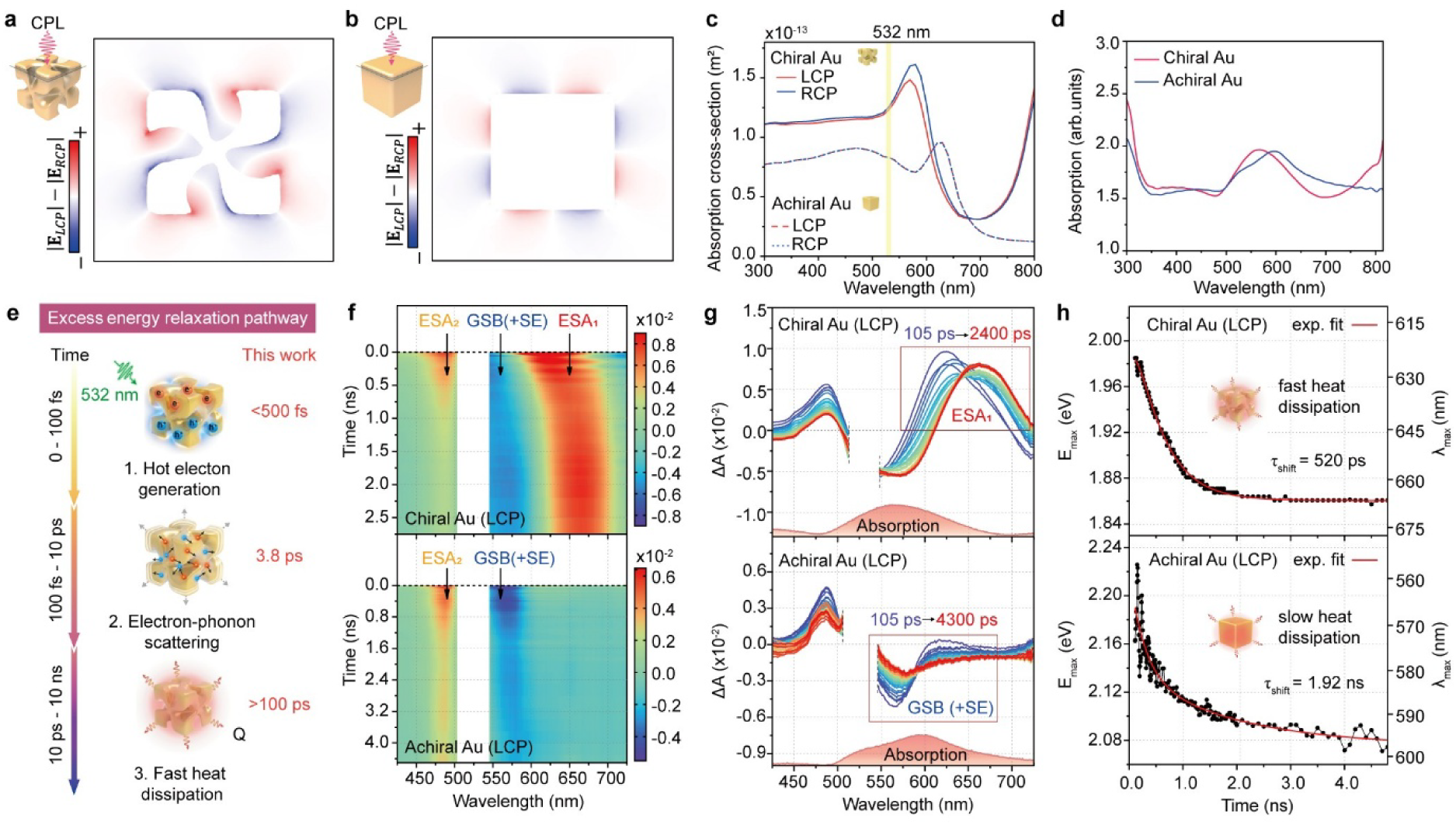
CPL-induced plasmonic energy localization and heat dissipation in chiral Au nanostructures. (a,b) Numerically simulated differential electric-field maps (|*E_LCP_*| − |*E_RCP_*|), and (c) simulated absorption cross-sections for chiral and achiral AuNPs under CPL illumination. The chiral Au exhibits a pronounced asymmetric near-field distribution, whereas the achiral Au shows a comparatively symmetric response, highlighting the moderate polarization-dependent optical response of the chiral structure and the relatively weak polarization sensitivity of the achiral counterpart. (d) Experimental absorption spectra of chiral and achiral AuNPs in water measured using a conventional visible light source. Note that the absorption cross-section of chiral AuNPs is higher than that of achiral AuNPs at 532 nm. (e) Schematic illustration of the plasmon-mediated energy relaxation processes of an AuNP under femtosecond 532 nm excitation. (f, g) Transient absorption (TA) contour maps under LCP excitation and representative TA spectra of chiral and achiral AuNPs (150 ps intervals) after ultrafast electron-phonon coupling. ESA, GSB, and SE denote excited-state absorption, ground-state bleach, and stimulated emission, respectively. For clarity, the strong pump scattering at 532 nm was removed (vertical white box) (h) First-moment analysis (intensity-weighted center position, E_max_ in eV, or λ_max_ in nm) for the chiral (ESA_1_ band) and achiral (GSB band) AuNPs.

Next, we systematically investigated the sequential energy-relaxation pathways of the AuNPs dispersed in water using femtosecond transient absorption (TA) spectroscopy under 532 nm excitation with varying CPL direction (**Fig. 2e-h, and Extended Data Fig. 3a**). In the TA spectra, the chiral AuNPs exhibited two distinct excited state absorption bands (ESA_1_, ∼650 nm; ESA_2_, ∼480 nm) and one ground state bleach (GSB, ∼580 nm), whereas the achiral AuNPs showed relatively simple spectral features with negligible ESA_1_ around 650 nm (**Extended Data Fig. 3b-c**). In both samples, the contribution from stimulated emission (SE), corresponding to plasmon-mediated photoluminescence (PL)^47^, was not spectrally resolved because of the negligible Stokes shifts of the PL bands (**Extended Data Fig. 4a-b**).

In the TA spectra, the ESA_2_ band exhibited negligible dynamic spectral shifts in both samples, allowing the TA kinetics at 490 nm to directly reflect the population relaxation of the photoexcited plasmon states. Both samples showed comparable relaxation dynamics (**Extended Data Fig. 3d-e**), consistent with the well-established relaxation scheme for AuNPs illustrated in **Fig. 2e**^48^. Importantly, only the chiral AuNPs exhibited a clear CPL dependence in the TA amplitude, which was substantially enhanced under chirality-matched LCP excitation^49^. These results provide direct evidence for chirality-driven photonic PCR. Moreover, although the ultrafast electron-phonon coupling (3.7-3.8 ps) was nearly insensitive to the chiral morphology, the subsequent population relaxation processes (>100 ps) were accelerated in the chiral AuNPs. This suggests that system-bath interactions are enhanced in the chiral AuNPs, leading to faster deactivation. On the other hand, coherent lattice vibrations of the cubic AuNPs, induced by rapid lattice heating through the electron-phonon coupling, were also resolved in the TA spectra above 550 nm (**Extended Data Fig. 4c-d**)^50,51^. Linear-prediction singular value decomposition (LPSVD) analysis resolved two damped oscillations^52^, whose frequencies decreased upon introducing the chiral morphology. The reduced frequencies may arise from both the unique hollow structure and the relatively high lattice temperature of the chiral AuNPs^48,53^.

In this work, to quantitatively probe the slow heat-dissipation process of AuNPs in solution, we examined the dynamic spectral shifts of their plasmon-mediated bands following the ultrafast electron-phonon coupling (<10 ps) (**Fig. 2f-h**). For both samples, the GSB bands were initially blue-shifted relative to the corresponding stationary plasmon absorptions (red-shaded curves in **Fig. 2g**) and gradually shifted toward their equilibrium positions on a 2∼4 ns timescale. The strong ESA_1_ band of the chiral AuNPs exhibited a similar trend. Intensity-weighted spectral analysis (1^st^ moment, **Fig. 2h**) further showed that the chiral AuNPs underwent a pronounced and relatively fast bathochromic shift (τ_shift_=520 ps), whereas the achiral AuNPs exhibited much weaker spectral evolution that persisted for up to 4 ns (τ_shift_=1.92 ns). Although the origin of the initially blue-shifted plasmon bands in the cube-shaped AuNPs remains unclear, the subsequent slow shifts of the TA bands toward their stationary positions likely reflect heat dissipation from the AuNPs and the surrounding water until thermal equilibrium is established. It is well known that the heat-dissipation rate increases as the size of AuNP decreases because of the larger surface-to-volume ratio^54^, indicating that the faster heat-dissipation and thermalization of the chiral AuNP can be attributed to its unique hollow structure and associated CPL-dependent field localization. The relatively strong bathochromic shift of the chiral AuNP may originate from a highly modulated dielectric response of the surrounding water and partial lattice deformation through intense local heating^55,56^.

On the other hand, rapid cooling of the PCR system during the dark stage of each PCR cycle is another key factor governing its performance. In this context, we examined whether non-activated chiral AuNPs can act as effective heat conductors in aqueous solution during the dark stage (**Extended Data Fig. 5**). The proposed fast-cooling reaction scheme is illustrated in **Extended Data Fig 5a**. Au nano-bipyramid (NBP) was employed as a spectroscopic probe because it has a longitudinal plasmon band at ∼750 nm^57^, which is well separated from the plasmon bands of the isotropic AuNPs (**Extended Data Fig 5b-d**). Accordingly, femtosecond excitation at 700 nm can selectively activate the NBPs, allowing quantitative monitoring of the dynamic spectral shifts of the GSB band around 750 nm in the presence of non-activated additives (chiral and achiral AuNPs). The absorbances of the chiral and achiral AuNPs were matched at 532 nm for the TA measurements. Thus, based on the simulation results (**Fig. 2c**), the particle density of chiral AuNPs is expected to be lower than that of achiral AuNPs. Moreover, the additive concentrations relative to the NBPs were kept sufficiently low that they produced negligible TA signals under 700 nm excitation (**Extended Data Fig. 5e**).

The pristine NBP solution exhibited well-separated ESA and GSB(+SE) bands around 600 nm and 750 nm, respectively. We then examined the slow dynamic spectral shift of the GSB band for three controlled samples in water: NBP, NBP+Chiral Au, and NBP+Achiral Au (**Extended Data Fig 5f-j**). Global fitting analysis with two log-normal functions was used to account for the spectral overlap between the ESA and GSB bands near 700 nm as well as their asymmetric Gaussian shapes; this analysis provided the raw TA spectra together with the fitted curves at representative time-delays and the corresponding fitting parameters for the deconvoluted GSB band, including the center position shift, τ_shift_ and the peak amplitude, *h*(t). As expected, the dynamic spectral shift accelerated markedly in the presence of chiral AuNPs (*τ*_shift_=740 ps → 454 ps), accompanied by a reduced spectral shift (*λ*_base_=786.8 nm → 762.6 nm) in the plateau region beyond 1 ns. These changes likely reflect enhanced heat dissipation in the local environment surrounding the photoexcited NBP. The corresponding population relaxation extracted from the temporal evolution of the peak amplitude also accelerated substantially (*τ*_2_=2080 ps → 865 ps), further supporting enhanced system-bath coupling. The mixture containing achiral AuNPs also showed similar deviations, but their influence on GSB dynamics was considerably weaker (τ_shift_=825 ps, λ_base_=771.6 nm, and τ_2_=1470 ps), even though the effective particle density of achiral AuNPs is expected to be higher than that of chiral AuNPs.

In summary, we performed two complementary TA experiments to establish the mechanistic foundation for rapid and polarization-sensitive thermal cycling used in chirality-driven photonic PCR. The overall results show that upon 532 nm irradiation, chiral AuNPs exhibit stronger CPL-dependent field localization, enhanced absorption near the excitation wavelength, and faster heat dissipation than achiral AuNPs. We further found that during the dark stage, an aqueous solution containing non-activated chiral AuNPs can act as an effective heat reservoir by facilitating thermal transport.

### Rapid and polarization-selective photothermal cycling

We next evaluated whether the photophysical advantages of chiral AuNPs translate into practical photothermal cycling performance under PCR-relevant conditions. The design principle is that chiral plasmonic nanoparticles, owing to their asymmetric interaction with CPL, generate stronger photothermal output than achiral particles under otherwise matched irradiation conditions (**Fig. 3a**). Experimental CD spectra confirmed that only the chiral AuNPs exhibited a pronounced chiroptical response overlapping the optical excitation window, whereas the achiral AuNPs showed a near-featureless CD signal; this experimental asymmetry was well reproduced by the simulated circular differential extinction spectrum of the chiral AuNPs (**Fig. 3b**). Simulated extinction cross-sections further showed that the chiral AuNPs responded differently to LCP and RCP illumination, whereas the achiral AuNPs showed nearly identical extinction under both polarizations (**Fig. 3d**), indicating that polarization-selective light absorption is a structural property specific to the chiral geometry.

**Figure 3.**
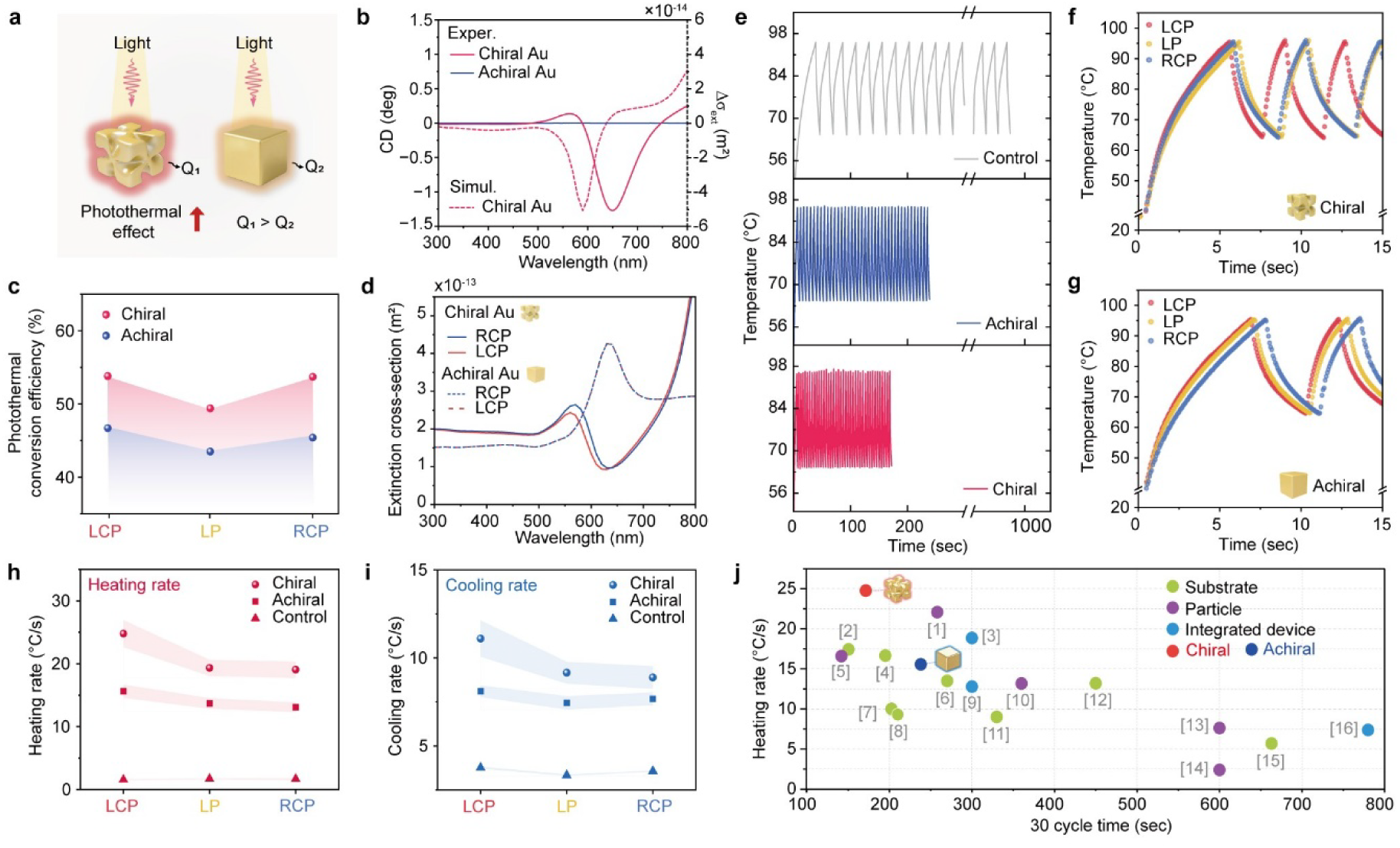
Photothermal performance of chiral and achiral AuNPs under CPL. (a) Schematic illustration of the photothermal effect in chiral and achiral AuNPs under light irradiation. Q denotes the heat generated by each nanostructure, with the chiral AuNP producing more heat (Q) than the achiral AuNP (Q_1_ > Q_2_). (b) Experimental CD spectra of chiral and achiral AuNPs and simulated circular differential extinction cross-section (Δ*σ_ext_*) spectrum of chiral AuNPs. (c) Photothermal conversion efficiencies of chiral and achiral AuNPs under LCP, LP, and RCP irradiation, showing enhanced photothermal performance for the chiral structures. (d) Simulated extinction cross-section of chiral and achiral AuNPs under RCP and LCP excitation. (e) Temperature cycling profiles recorded over 40 heating–cooling cycles for deionized water without nanoparticles (control), achiral AuNPs and chiral AuNPs under LCP irradiation. (f,g) Representative heating and cooling curves of (f) chiral AuNPs and (g) achiral AuNPs recorded over a 15 s interval during the photothermal cycling experiments under LCP, LP, and RCP irradiation. (h,i) Comparison of (h) heating rates and (i) cooling rates of deionized water (control), achiral AuNPs, and chiral AuNPs under LCP, LP, and RCP irradiation. Shaded regions indicate the standard deviation from *n* = 3 independent measurements. (j) Comparison of photothermal heating rates as a function of 30-cycle amplification time, highlighting the performance of the present chiral AuNP system relative to previously reported plasmonic photothermal platforms. Numbers in brackets correspond to the references listed in Supplementary Table 2.

To determine whether this optical asymmetry translates into functional photothermal output, we measured the photothermal conversion efficiency of both particle types under LCP, linearly polarized (LP), and RCP irradiation. Chiral AuNPs consistently outperformed achiral AuNPs across all three polarization states (**Fig. 3c**), confirming that the enhanced near-field localization and polarization-selective absorption identified above directly translate into more efficient photothermal conversion.

We then evaluated temperature-cycling performance under repeated optical irradiation across the full PCR-relevant range. Notably, chiral AuNPs reached the PCR temperature window more rapidly and maintained repeated thermal oscillations over 40 cycles within a shorter time frame than achiral AuNPs (**Fig. 3e**). This enhanced cycling performance was consistent across polarization states, as further supported by additional real-time cycling profiles recorded under LP and RCP irradiation (**Extended Data Fig. 6**). Representative short-time temperature traces further resolved that chiral AuNPs exhibited rapid thermal oscillations under all three polarizations, with the fastest response observed under LCP (**Fig. 3f**), whereas achiral AuNPs showed consistently slower heating and cooling under the same conditions (**Fig. 3g**).

Quantitative analysis confirmed that chiral AuNPs achieved higher heating rates than achiral AuNPs and the nanoparticle-free control under all polarization states (**Fig. 3h**). The cooling rates followed the same trend, indicating that structural chirality improves not only light-induced heating but also thermal recovery during the cooling step required for PCR cycling (**Fig. 3i**). Optimization experiments further defined the operating window for photonic PCR by varying nanoparticle optical density, laser power and reaction volume, and control experiments at 655 nm confirmed that the chiral Au system retained superior photothermal performance and supported successful DNA amplification under alternative excitation conditions (**Extended Data Fig. 7**).

Finally, we benchmarked the present chiral AuNP platform against previously reported photothermal PCR systems. The chiral Au system occupied a favorable performance regime, combining high heating rates with shortened 30-cycle amplification times relative to substrate-, particle- and integrated-device-based photothermal PCR platforms reported in the literature (**Fig. 3j**). Together, these results demonstrate that chiral AuNPs act as efficient and polarization-selective photothermal transducers, enabling rapid and stable heating–cooling cycles suitable for accelerated photonic PCR.

### Chiral near-field-induced conformational activation of *Taq* DNA polymerase

To elucidate the biochemical basis for the accelerated amplification rate, we investigated the interaction between strongly localized chiral electromagnetic fields and the conformational dynamics of Taq DNA polymerase during DNA synthesis (**Fig. 4**). Taq polymerase catalysis proceeds through a series of conformational transitions: the enzyme moves from the apo form (1KTQ) to a DNA-bound open binary complex (4KTQ), to an open ternary complex (2KTQ) upon dNTP binding, and finally to a closed ternary complex (3KTQ) that supports nucleotide incorporation (**Fig. 4a**). This final, rate-limiting step requires a conformational “O-helix closure” transition to establish the closed ternary state, followed by the release of pyrophosphate and the reopening of the enzyme to proceed to the next catalytic cycle. Structural comparison of the 4KTQ and the 3KTQ showed that the distance between Leu544 and Asp655 decreased from 39.5 Å in the open state to 27.7 Å in the closed ternary state, a displacement of 11.8 Å (**Fig. 4b,c**). This large change in distance suggests substantial structural rearrangements near the polymerase active site. Analysis of residue-level Cα displacements revealed that the most prominent structural changes were concentrated in the palm C-terminal region (residues 640–660, maximum 15.4 Å) and the O-helical domain (residues 520–545), while regions outside the simulation area showed displacements of less than 3 Å, indicating no structural changes during the transition process (**Fig. 4d**).

**Figure 4.**
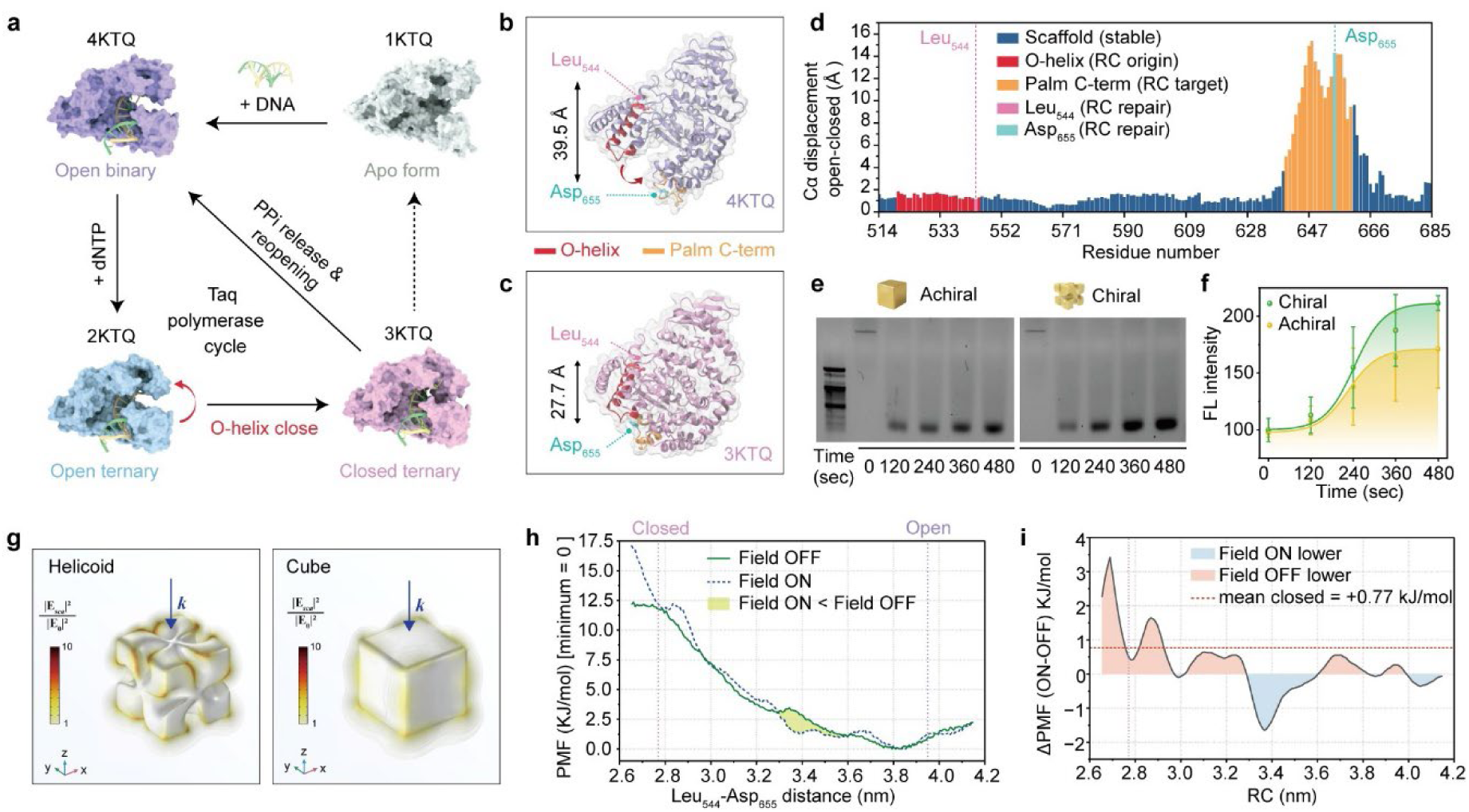
Enhanced photonic PCR through conformational activation of Taq polymerase. (a) Schematic representation of the *Taq* polymerase catalytic cycle based on representative crystal structures: apo form (1KTQ), DNA-bound open binary complex (4KTQ), open ternary complex after dNTP binding (2KTQ), and closed ternary complex (3KTQ). The transition from the open to closed ternary state involves O-helix closure, followed by PPi release and reopening for the next catalytic cycle. (b,c) Structural comparison of *Taq* polymerase in the (b) open binary state and (c) closed active ternary state, showing the decrease in Leu_544_–Asp_655_ distance from 39.5 Å to 27.7 Å during conformational closure. (d) Residue-level Cα displacement analysis during the open-to-closed transition, highlighting large structural rearrangements in the O-helix region and palm C-terminal region. (e) Agarose gel electrophoresis analysis of photonic PCR products amplified on chiral and achiral AuNPs substrates under LCP irradiation at different reaction times. (f) Time-dependent fluorescence intensity profiles of PCR amplification on chiral and achiral AuNPs under LCP irradiation. Error bars represent s.d. from *n* = 3 independent measurements for each condition. (g) Simulated electromagnetic field distributions around chiral helicoid and achiral cubic, showing gap-localized fields for the helicoid and more extended surrounding fields for the cube. (h) Potential of mean force (PMF) profiles along the Leu_544_–Asp_655_ reaction coordinate with and without the numerically derived chiral electromagnetic field. (i) Difference in PMF between field-on and field-off states, showing the field-dependent energetic modulation of polymerase conformational dynamics.

We hypothesized that the strongly asymmetric near-fields generated by chiral nanostructures couple favorably to these local structural dipoles, modulating the free-energy landscape of O-helix closure. To test this, we calculated the electromagnetic field distributions around chiral helical nanostructures and achiral cubic nanostructures (**Fig. 4g**). The helicoidal Au nanostructures generated a field localized to the gap, whereas the achiral cubic nanostructures generated a more broadly distributed electromagnetic field around the particle surface. To directly evaluate the energetic effects of this electromagnetic field asymmetry, we performed umbrella sampling molecular dynamics simulations using the Leu544–Asp655 Cα distance as the response coordinate and compared a system with and without the application of a chiral electromagnetic field retrieved from FEM simulation (LCP asymmetry, *r*=3 nm at the nanoparticle surface) (**Extended Data Fig. 8a-c, f)**. Mean Force Potential (PMF) analysis showed that the chiral field makes the closed O-helix slightly less stable than the open state (ΔΔG = 0.54 kJ/mol). This is consistent with the fact that the energy barrier required for O-helix opening is lowered by the chiral field (**Fig. 4h, i**). Furthermore, re-equilibration simulations under progressively stronger near-field conditions (reduced probe radius, *r*=1.5– 2.0 nm) further showed that all 15 sampling windows underwent substantially larger structural deviations toward the open state than the ensemble average across all windows, providing direct evidence that the chiral near-field preferentially drives closed-state conformations toward the open form (**Extended Data Fig. 8d-e**). These simulation results support the possibility that the chiral near-field can act directly on polymerase conformational dynamics in addition to photothermal effects.

To experimentally assess this hypothesis, we monitored amplification kinetics using time-dependent agarose gel electrophoresis and fluorescence measurements under controlled polarization states. Under LCP irradiation, chiral AuNP-assisted photonic PCR produced detectable DNA products earlier and with stronger band intensities than the achiral AuNP system, accompanied by a markedly faster increase in fluorescence intensity over the same time course (**Fig. 4e,f**). Under RCP irradiation, both systems supported DNA amplification and time-dependent fluorescence growth, but the enhancement associated with the chiral Au system was less pronounced than under LCP excitation (**Extended Data Fig 8g-h**). Together, the structural analysis, electromagnetic simulations, gel electrophoresis and fluorescence measurements suggest that chiral plasmonic nanoparticles do more than function as passive nanoscale heaters. Instead, the coupling between structural chirality and optical chirality may create a favorable biophysical environment for polymerase conformational transitions, thereby accelerating DNA amplification during photonic PCR.

### Label-free chiroptical detection of SARS-CoV-2

We finally evaluated whether the chirality-driven photonic PCR platform could be extended to a diagnostically relevant viral target and combined with label-free chiroptical readout. As a model target, we selected a 113-bp region of the SARS-CoV-2 envelope (E) gene encoded in a plasmid template, representing a defined viral sequence within the approximately 30-kb SARS-CoV-2 genome (**Fig. 5a**). This target allowed us to directly compare conventional thermal cycling, achiral and chiral AuNP-assisted photonic PCR under controlled amplification conditions. Temperature profiles showed that the conventional PCR machine required substantially longer cycling times than either photonic PCR systems, whereas AuNP-mediated optical heating enabled rapid repeated thermal oscillations (**Fig. 5b**). Among the photonic system, chiral AuNPs completed 30 thermal cycles faster than achiral AuNPs under the same polarization conditions, with the shortest cycling time achieved under LCP irradiation (**Fig. 5c**). These results extend the polarization-selective photothermal advantage of the chiral Au platform, established above for model DNA templates, to a clinically relevant viral amplification workflow. Additional gel electrophoresis analyses confirmed that the photonic PCR system amplified DNA across different nanoparticle concentrations, cycle numbers and polarization states, while conventional PCR served as a benchmark control (**Extended Data Fig. 9**).

**Figure 5.**
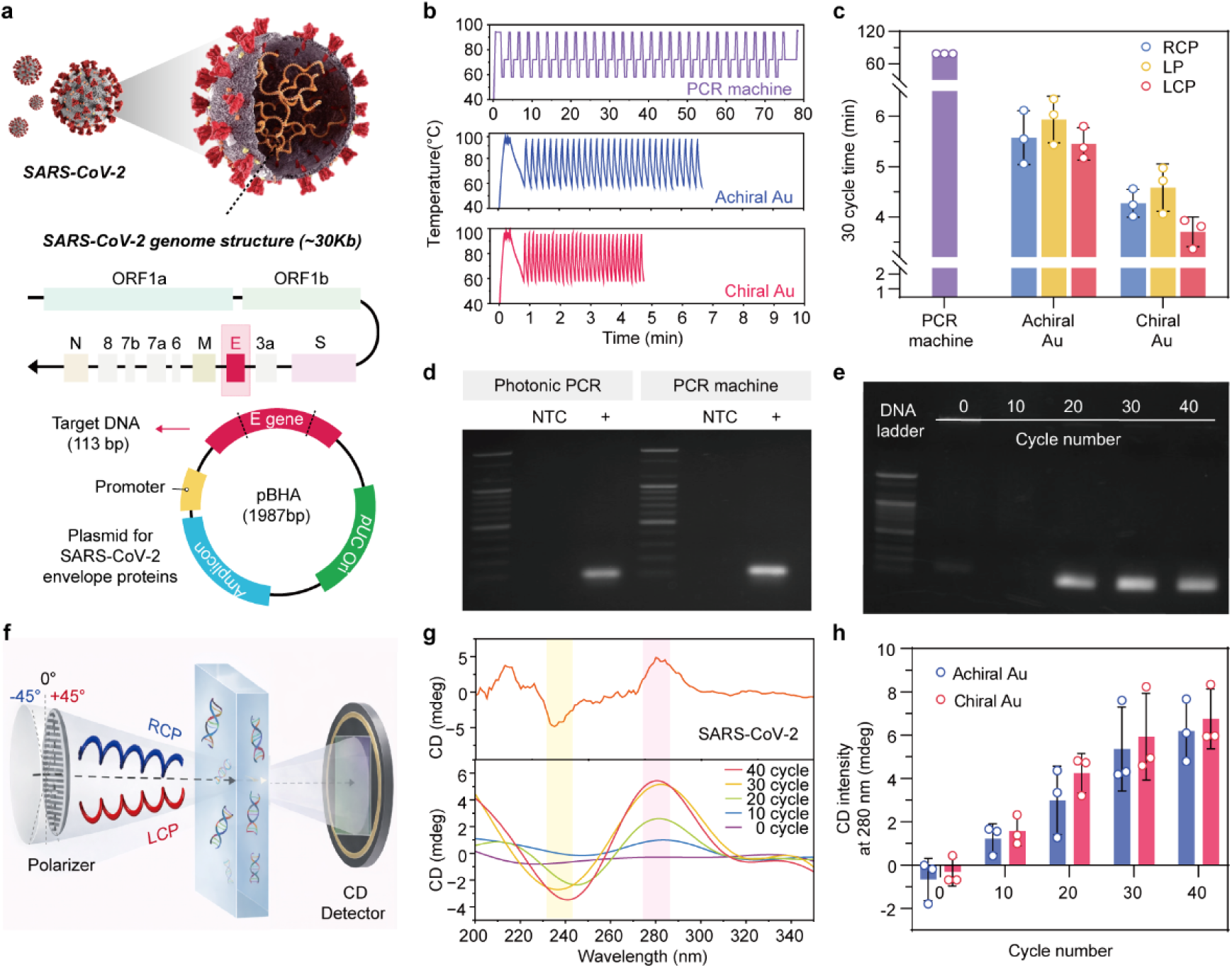
CD analysis of the photonic PCR product of SARS-CoV-2. (a) Schematic illustration of the SARS-CoV-2 genome (∼30 kb) and plasmid construction containing the envelope (*E*) gene target used for PCR amplification (113 bp). (b) Representative temperature profiles during PCR cycling using a conventional PCR machine, achiral AuNPs, and chiral AuNPs. (c) Comparison of the total reaction time for 30 PCR cycles using a conventional PCR machine with achiral AuNPs and chiral AuNPs under different polarization conditions (LCP, LP, and RCP). Error bars represent s.d. from *n* = 3 independent measurements for each condition. (d) Agarose gel electrophoresis analysis of SARS-CoV-2 E gene amplicons generated by photonic PCR and conventional PCR. NTC denotes the no-template control. (e) Cycle-dependent gel electrophoresis analysis showing progressive accumulation of PCR products from 0 to 40 cycles. (f) Schematic of the CD measurement setup, in which LCP and RCP components are transmitted through the PCR product solution and detected by a CD detector. (g) CD spectra of SARS-CoV-2 DNA amplicons measured after different PCR cycle numbers, showing progressive development of the chiroptical signal without fluorescent labelling. (h) Quantitative comparison of CD intensity at 280 nm as a function of PCR cycle number for achiral and chiral AuNP-assisted PCR, highlighting enhanced signal amplification in the chiral system. Error bars represent s.d. from *n* = 3 independent measurements for each condition.

We next verified the amplified SARS-CoV-2 target products by agarose gel electrophoresis. Chiral AuNP-assisted photonic PCR produced a clear target band comparable to that obtained using a conventional PCR machine, whereas no-template controls showed no detectable amplification in either system (**Fig. 5d**). Cycle-dependent gel analysis further showed progressive accumulation of the target product from 0 to 40 cycles, confirming that the optical cycling protocol supports stepwise DNA amplification rather than nonspecific thermal artifacts (**Fig. 5e**). Together, these results establish that rapid chiral photonic cycling produces sequence-specific viral amplicons under compressed reaction time.

To integrate amplification with label-free detection, we monitored the intrinsic CD response of the amplified DNA products directly, without any fluorescent label. In this configuration, incident light was resolved into left- and right-handed circularly polarized components before passing through the PCR product solution, and the resulting differential absorption was recorded by a CD detector (**Fig. 5f**). Because double-stranded DNA adopts a chiral helical structure, the accumulation of PCR products is expected to increase the ultraviolet CD response associated with the DNA backbone and base-stacking. Consistently, CD spectra of SARS-CoV-2 amplicons showed cycle-dependent signal development, with increasingly pronounced chiroptical features as the number of PCR cycles increased from 0 to 40 (**Fig. 5g**). Quantitative analysis at 280 nm further showed that CD intensity increased with cycle number, with the chiral AuNP-assisted PCR producing a stronger chiroptical signal than the achiral system at later amplification cycles (**Fig. 5h**). We further validated the generality of this CD-based readout using lambda DNA amplification products. The amplified DNA displayed the characteristic B-form CD signature, including a negative band near 245 nm and a positive band near 275 nm, with CD intensity increasing with both DNA concentration and PCR cycle number (**Extended Data Fig. 10a-h**). Control experiments in which chiral or achiral AuNPs were included during amplification using a conventional PCR machine also showed successful DNA amplification by gel electrophoresis and confirmed that the post-amplification CD signals were enhanced relative to the pre-amplification state (**Extended Data Fig. 10i-l**). These results confirm that the observed CD response originates from amplified DNA products and can serve as a dye-free optical readout of PCR amplification. Together, these data demonstrate that chiral AuNP-assisted photonic PCR enables rapid amplification of a SARS-CoV-2 gene target and direct label-free detection of the resulting DNA products through their native chiroptical response.

## Conclusions

In conclusion, we establish a paradigm shift in molecular diagnostics by developing a chirality-driven photonic PCR platform that enables both ultra-rapid genetic amplification and direct, label-free chiroptical detection. Chiral AuNPs exhibited polarization-dependent excitation and markedly accelerated energy dissipation than achiral AuNPs, with the spectral relaxation time associated with thermal equilibration decreasing from 1.92 ns to 520 ps, as resolved by femtosecond TA spectroscopy. These photophysical advantages support faster heating and cooling, enabling amplification of a plasmid-encoded SARS-CoV-2 E-gene target over 30 thermal cycles in less than 5 min. In addition, localized chiral near-fields can modulate the conformational dynamics of Taq DNA polymerase by favoring O-helix reopening, providing apotential contribution to rapid amplification. Beyond rapid amplification, we show that the resulting DNA products can be detected directly through their intrinsic CD response, eliminating the need for fluorescent dyes or labelled probes. Validated on both a model genomic template and a SARS-CoV-2 target under practical reaction timelines, our platform provides a highly versatile, universally adaptable architectural blueprint for next-generation point-of-care testing devices.

## Methods

### Chemicals and reagents

Gold (III) chloride trihydrate (HAuCl_4_.3H_2_O; ≥99.9% trace metals basis), L-Ascorbic acid (AA, 99%), sodium borohydride (NaBH_4_, 99%), Potassium iodide (KI ≥99%), L-glutathione (L-GSH, 98%), hexadecyltrimethylammonium bromide (CTAB, 99%), cetyltrimethylammonium chloride solution (CTAC, 25 wt% in H_2_O) were purchased from Sigma Aldrich.

Reagents for conventional PCR amplification, including 10 mM dNTPs, 10× reaction buffer (containing MgCl₂), Taq DNA polymerase, gene-specific forward and reverse primers, agarose, 50× Tris-acetate-EDTA (TAE) buffer, 6× agarose gel loading buffer and a 100-bp DNA ladder were all purchased from Bioneer (Daejeon, South Korea). Lambda (λ) DNA was purchased from Roche Applied Science (250 μg/ml), and a 113-bp fragment of the SARS-CoV-2 envelope (E) gene, together with the corresponding forward and reverse primers, was obtained from Bioneer. Mineral oil was purchased from Sigma-Aldrich. SYBR™ Green I Nucleic Acid Gel Stain were purchased from Thermofisher Scientific. RNase-free water was purchased from Welgene. All aqueous solutions were prepared using high-purity deionized (DI)-water (18.2 MΩ cm).

### Characterization

Scanning electron microscopy (SEM) images were obtained by an energy-dispersive Schottky-field emission scanning electron microscope (JSM-7610). CD spectra were acquired using the JASCO J-1500 CD spectrometer. UV–vis absorption spectra were measured using the Cary 5000 (Varian) spectrometers. Gel images were taken using ImageQuant 800 and gel documentation system (FLB-001BCM, Cellgentek). Conventional PCR was conducted by Applied Biosystems (Step One Plus Real Time PCR System).

### Synthesis of chiral and achiral gold nanoparticles

Chiral gold nanoparticles were synthesized via the peptide-mediated growth method^38,39^. Specifically, the spherical seeds were first synthesized by adding NaBH_4_ (10 mM, 0.8 mL) into a HAuCl_4_ (10 mM, 0.25 mL) containing CTAB (100 mM, 7.5 mL) solution and the solution was kept at 28 °C for 3 h to decompose the remaining NaBH_4_. Then the octahedral seeds were prepared by growing the spherical seeds in the growth solution, which is composed of 8 ml of water, HAuCl_4_ (10 mM, 0.2 mL), CTAB (100 mM, 1.6 mL), and AA (50 mM, 0.95 mL). After 15 min incubation in 30 °C water bath, the solutions were centrifuged (6708 g, 60 s) and redispersed in 1 mM CTAB solution. Finally, the octahedral seeds were injected in the growth solution prepared by mixing CTAB (100 mM, 0.8 ml), HAuCl_4_ (10 mM, 0.2 ml), AA (100 mM, 0.475 ml) and L-GSH (10 mM, 2.75 µl), sequentially. For the achiral nanoparticles, no chiral ligand was added. After the seeds were added, the reaction solution was left for 2 h at 30 °C water bath without disturbing. The color was changed from transparent to blue with large scattering. Finally, the solution was centrifuged twice (1,677 g, 60 s) to eliminate unreacted reagents and was re-dispersed in a 1 mM CTAB solution for further use.

### Photonic PCR thermal cycler setup and control system

A custom-built photonic PCR thermal cycling platform was constructed on an optical table to ensure stable laser alignment and reproducible operation throughout the experiments. Photothermal heating was induced by CPL laser irradiation. To enhance light utilization and photothermal conversion efficiency, transmitted laser light was reflected back toward the reaction tube using a reflective aluminum foil layer wrapped around the lower section of the tube. An optical shutter positioned between the laser source and the sample regulated laser exposure according to the measured sample temperature. The shutter was fabricated using a thin stainless-steel plate coupled to a 180° servo motor. The reaction temperature was continuously monitored using a thermocouple inserted into the sample solution. Temperature data were acquired in real time using a Keithley 2000 multimeter and processed by a computer-based control program that regulated shutter operation according to user-defined temperature limits. To accelerate cooling, a 120 mm 12 V DC cooling fan was activated when the shutter was closed and deactivated when the shutter was opened. The shutter and cooling fan were controlled through an Arduino Uno board with the control program. A control program was developed to perform automated thermal cycling based on real-time temperature feedback. The program included pre-heating and temperature-hold functions for PCR operation. Detailed photographs of the experimental setup, conceptual diagram and control logic flow are provided in Extended Data Fig. 2.

### Photonic PCR Mixture Preparation

The photonic PCR reaction mixture was prepared in a total volume of 10 μl, consisting of 1× reaction buffer (10 mM Tris-HCl, 50 mM KCl and 1.5 mM MgCl_2_), 0.2 mM each dNTP (Bioneer), 1.5 U Taq DNA polymerase (Bioneer), 2.5 μM each of the forward and reverse primers (Bioneer; see Supplementary Table 3), 0.5 μl of template DNA (λDNA, Roche Applied Science; or the SARS-CoV-2 plasmid template, Bioneer; see Supplementary Table 3), and chiral or achiral AuNPs (5 μl per reaction; optical density of 20 at 532 nm), with nuclease-free water added to bring the mixture to its final volume. The mixture was loaded into a PCR tube (Thermo Fisher Scientific) and overlaid with 10 μl of mineral oil to prevent evaporation, and the tube was then subjected to thermal cycling in the custom-built photonic PCR device under CPL irradiation to drive amplification. Amplification was monitored by agarose gel electrophoresis and by the intrinsic CD signal of the amplified DNA product.

### Gel Electrophoresis

The products of photonic PCR were separated by electrophoresis at 135 V on a dry precast 2% agarose gel containing 1X SYBR safe (Invitrogen) as a staining dye for 15 min. The lengths of the products were determined relative to a 100 bp DNA Ladder (Invitrogen). Gel images were acquired using a gel documentation system (FLB-001BCM, Cellgentek) and analyzed using NEOimage for FluoroBox software.

### Time-resolved optical spectroscopy

We used a commercial Ti:sapphire regenerative amplifier system (Spectra Physics, Spitfire Ace) to generate amplified femtosecond pulses (> 1 mJ, 800 nm, and 1 kHz). We directed the 800 nm output to an optical parametric amplifier (OPA) to generate pump beams at 532 and 700 nm. We used a commercial TA spectroscopy system (Ultrafast Systems, Helios) to monitor the excited transient species. For the TA measurements, the excitation pulse energy was adjusted to about 250 nJ using a neutral density filter to avoid photodamage. The residual 800 nm pulse passed through an optical delay line and was focused into a 1 mm water cell to generate a white-light continuum, which was directed through the sample cell (2.0 mm optical path) and detected with a CCD detector. We controlled the pump-beam polarization state using an achromatic quarter-wave plate. The instrument response function (IRF) of the TA setup had a width of about 500 fs. For time-resolved photoluminescence (TRPL) measurements, we used the 532 nm OPA output to excite the samples and recorded the TRPL spectra in a 50 ns time window using a streak scope (Hamamatsu Photonics, C10627). The IRF width of the TRPL setup was about 300 ps. To obtain reliable PL spectral shapes, time-gated PL detection with a 590 nm long-pass filter was employed to examine the red-shoulder of the PL bands while suppressing the strong pump and solvent Raman scattering near time-zero.

### MD simulations for chiral field-based polymerase activity

The open and closed forms of the Taq protein were modeled based on the 4KTQ and 3KTQ crystal structures, respectively^58^. The A chain of 4KTQ was used as the initial structure. Deleted residues were substituted with alanine, and only functionally relevant regions (residues 514–685, a total of 172 residues), including the O-helix and palm C-terminal region, were used for the simulation. The structures were processed in GROMACS 2025.3 using the AMBER99SB-ILDN force field and the SPC/E water model^59–61^. The proteins were placed in a dodecahedral box, solvated with a minimum distance of 1.0 nm from the box edge, and then charged with Na^+^/Cl^−^ ions. The energy of the system was minimized using the steepest drop algorithm (tolerance 100 kJ/mol/nm). Subsequently, NVT equilibration was performed through a gradual temperature increase (100 ps) from 50 K to 345 K, during which the position of the protein backbone was restricted. Subsequently, NPT equilibration was performed first using a Berendsen pressure regulator (200 ps) followed by a Parrinello–Rahman pressure regulator (500 ps, τ_p = 2.0 ps, 1.0 bar) ^62,63^. All production simulations were performed at 345 K (V-rescale temperature regulator, τ_t = 0.1 ps) and 1.0 bar^64^. Electrostatic interactions were treated with PME (cutoff distance 1.2 nm), van der Waals interactions with a cutoff distance of 1.2 nm, and all bonds involving hydrogen atoms were constrained using the LINCS algorithm^65,66^. To generate initial structures encompassing open↔closed transitions, steering molecular dynamics (SMD) simulations were performed by pulling the Cα–Cα distance between Leu544 and Asp655 in the closed state direction at a velocity of −0.001 nm/ps and a force constant of 1000 kJ/mol/nm² at 4.117 nm^67,68^. Fifteen umbrella sampling windows were initialized by extracting frames at 0.1 nm intervals from valid pull trajectories (RC 2.70–4.10 nm). Each window was equilibrated (re-equilibrated) for 500 ps, after which a simulation was performed for 5 ns with a harmonic constraint (k = 1000 kJ/mol/nm²) centered on the target RC. We calculated the potential of mean force (PMF) using the weighted histogram analysis method.

We derived the local electric field experienced by the polymerase near a chiral helicoid AuNP under LCP excitation from wave optics simulations solved by COMSOL Multiphysics. The differential field intensity between LCP and RCP excitation at the helicoid gap (r = 3 nm from nanoparticle surface) was used to define the field asymmetry:

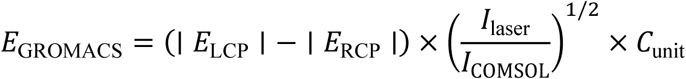

where ∣ *E*_LCP_ ∣ and ∣ *E*_RCP_ ∣ are the numerically computed optical nearfield magnitudes normalized to unit incident intensity (*|E_inc*| = 1 V/m), *I*_laser_/*I*_COMSOL_ is the laser-to-COMSOL intensity scaling factor, and *C*_unit_ converts V/m to V/nm. At the helicoid gap, the x-component differential was ∣ *E*_X_LCP_ ∣ − ∣ *E*_X_RCP_ ∣ = 1.158 − 1.429 = −0.271 V/m per unit incident field. After scaling to the experimental laser intensity, the applied field was (Ex, Ey, Ez) = (−1.2×10^− 5^, −9.0×10^−6^, −1.0×10^−6^) V/nm, consistent with LCP preferentially exciting the helicoid geometry (positive 532 nm CD). We applied this as a uniform external electric field in GROMACS during umbrella sampling windows for the field-ON condition. The field-OFF condition used identical simulation parameters without an applied field.

## Statistics and reproducibility

We present the data as mean ± standard deviation. In the SEM imaging and gel analysis experiments, we performed at least three independent experiments and showed representative images.

## Data availability

The data that supports the findings of this study are available from the corresponding author upon reasonable request.

## Acknowledgments

This research was supported by the Basic Science Research Program (Priority Research Institute) through the National Research Foundation (NRF) of Korea funded by the Ministry of Education (RS-2021-NR060138), by the National Research Lab 2.0 (NRL 2.0) (IMMS) through the NRF grant funded by the Korean government (MSIT and MOE) (RS-2025-16063688), and NRL 2.0 (K-BIGHEART) through the NRF grant funded by the Korean government (MSIT and MOE) (RS-2025-17492968). H. Kang acknowledges the financial support from the NRF grant funded by the Korean government (MSIT) (RS-2025-25420257). C. H. Kim acknowledges the financial support from the NRF grant (RS-2026-25483089). D. -I. Won acknowledges the financial support from the NRF grant funded by the Ministry of Education (No. RS-2024-00461946). K. T. Nam and J. H. Han appreciate the support from the National Research Foundation of Korea (NRF) (RS-2024-00409405).

## Author Contributions

L. P. Lee and D. H. Kim provided overall guidance and established the study framework. L. P. Lee conceived the original research concept of chiral photonic PCR and provided intellectual guidance throughout the project. C. H. Kim, K. T. Nam, D. Suh, J. W. Han and P. J. Yoo advised project strategy and provided experimental facilities. H. Kang conceived the concrete idea, carried out main experiments, analyzed data, and wrote the 1st manuscript. J. H. Han performed chiroptical simulations and Y. Kim conducted MD calculation. J. Park and D. Suh contributed to the design and set up for photonic PCR system. C. H. Kim, H. J. Lee and D. -I. Won conducted transient absorption spectroscopy measurements and analysis. Donghyun Kim, M. Kwon and Dabin Kim performed part of the main and supporting experiments. All authors revised the manuscript.

## Additional information

Supplementary Information is available for this paper. Correspondence and requests for materials should be addressed to XX. Reprints and permissions information is available at XX.

## Extended Data Legends

**Extended data figure 1.**
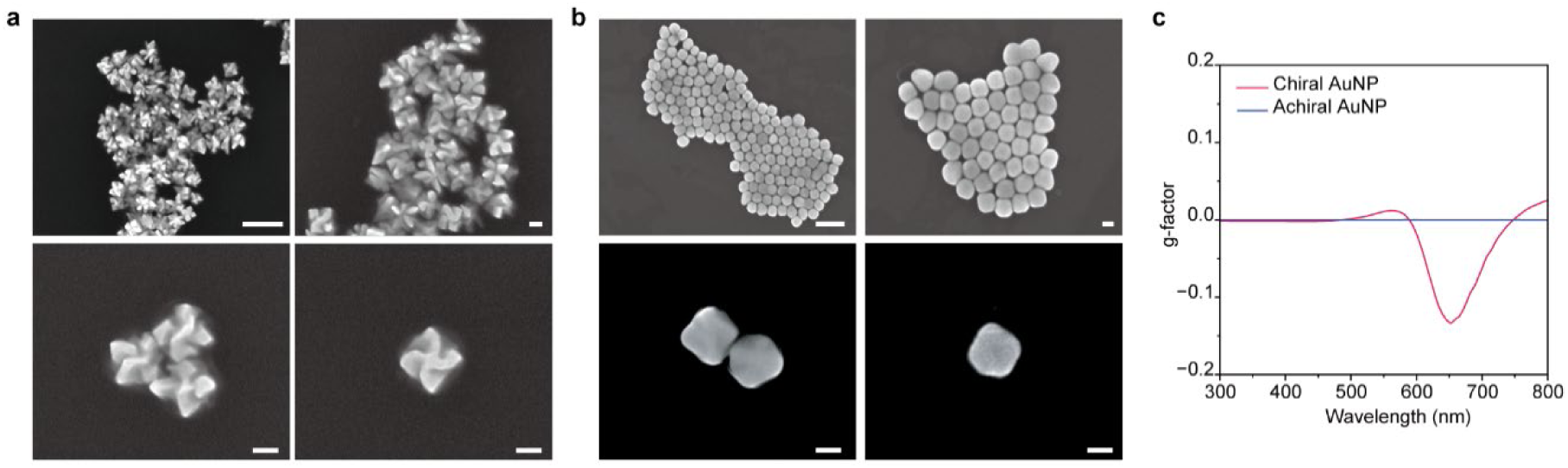
Morphological and chiroptical characterization of chiral and achiral AuNPs. (a) Representative SEM images of chiral AuNPs at different magnifications, showing the helicoidal morphology of the individual nanostructures. Scale bars, 500 nm (upper left) and 100 nm (upper right, lower left and lower right). (b) Representative SEM images of achiral AuNPs at different magnifications, showing individual achiral particle morphologies. Scale bars, 500 nm (upper left) and 100 nm (upper right, lower left and lower right). (c) g-factor spectra of chiral and achiral AuNPs. Chiral AuNPs exhibit a pronounced chiroptical response in the visible region, whereas achiral AuNPs show a near-zero g-factor over the measured wavelength range.

**Extended data figure 2.**
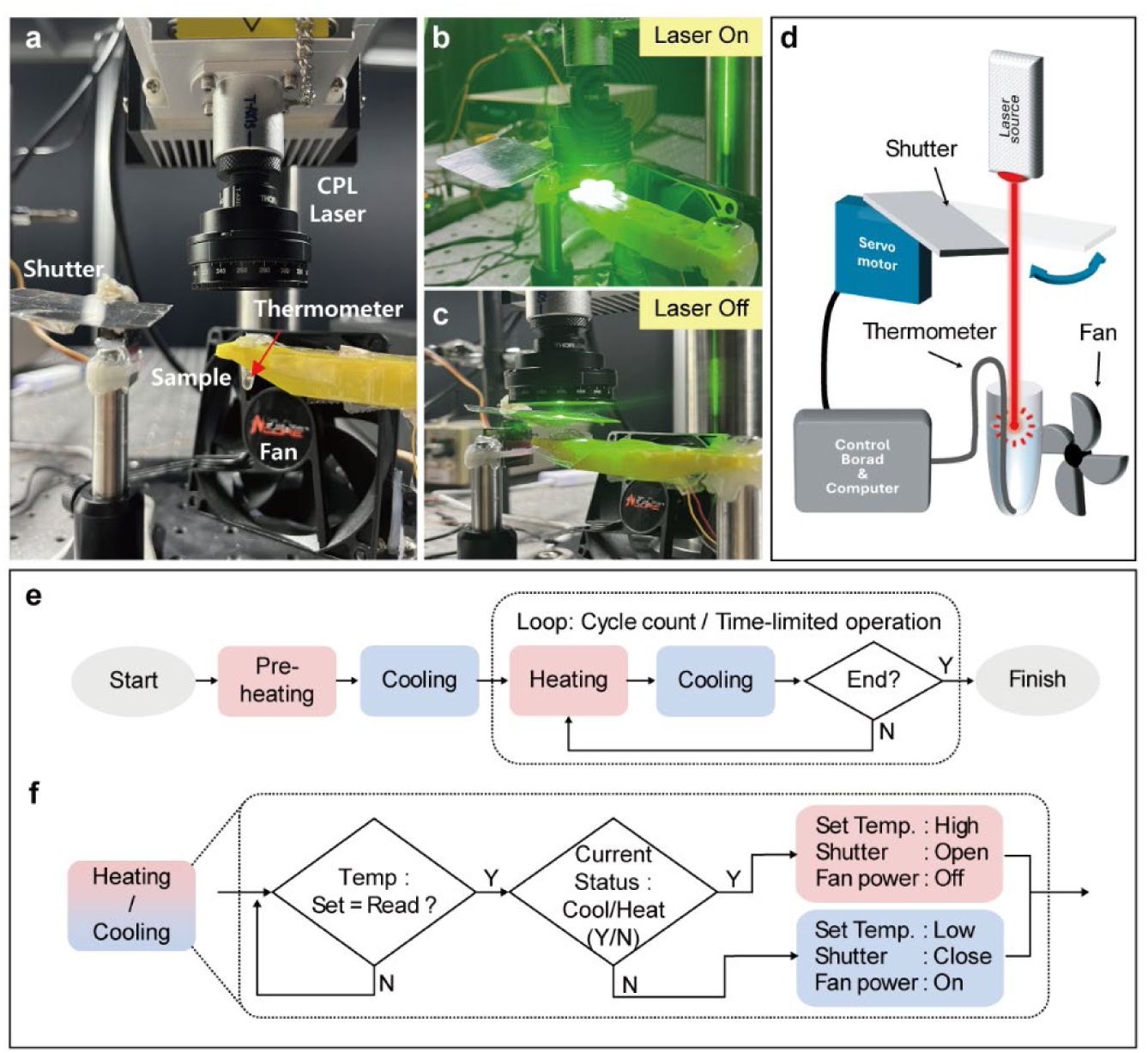
Plasmonic Photonic PCR device setup and thermal cycling control system. (a) Photograph of the custom-built photonic PCR system. The photothermal reaction stage consists of a CPL laser source (λ = 532 nm), optical shutter, sample stage, cooling fan and thermocouple, enabling real-time temperature monitoring and automated thermal cycling. (b,c) Photographs of the reaction stage under (b) laser-on and (c) laser-off conditions. During the heating step, the shutter is opened to allow the CPL laser to irradiate the sample and induce photothermal heating until the temperature reaches 95 °C. During the cooling step, the shutter is closed to block light irradiation, and the fan is activated to cool the sample until the temperature reaches 65 °C, after which the system switches to the next heating cycle. (d) Conceptual diagram of the photonic PCR control system, in which the optical shutter and cooling fan are regulated by a computer-connected control board according to the measured reaction temperature. (e) Logic flow of the CPL laser–based photonic PCR thermal cycling system. Thermal cycling was performed according to user-defined temperature limits, holding times and cycle number. For the PCR experiments in this work, the reaction mixture was pre-heated at 94–96 °C for polymerase activation before amplification. During PCR cycling, the optical shutter and cooling fan were automatically controlled using real-time temperature feedback to control sample heating and cooling. (f) Detailed logic flow for heating/cooling Control. When the measured temperature reaches the preset value, the system maintains that temperature for a defined period before automatically switching to the opposite thermal state. If the temperature does not reach the target value, the system continuously monitors the temperature until the condition is satisfied.

**Extended data figure 3.**
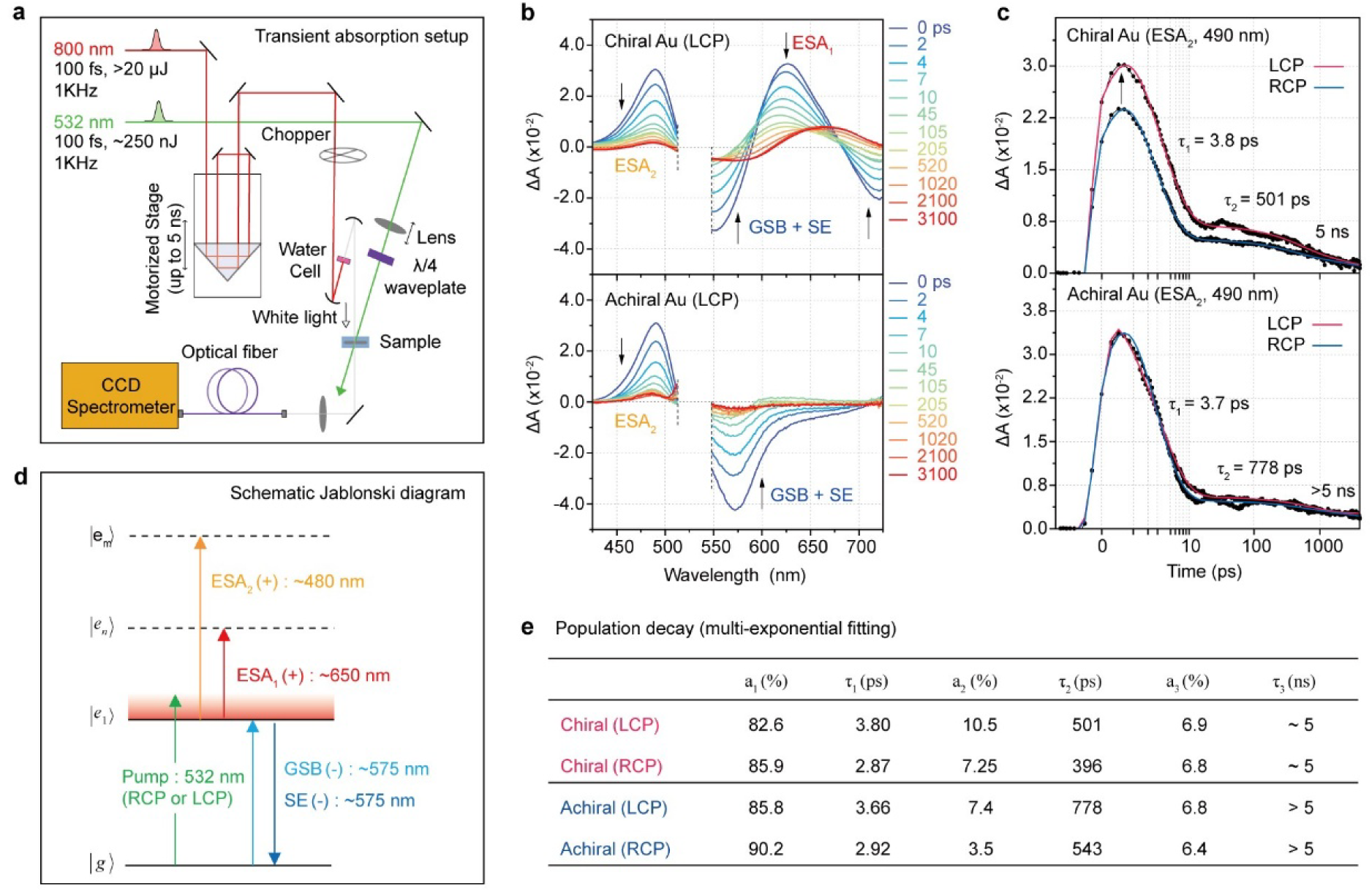
Femtosecond transient absorption (TA) characterization of chiral and achiral plasmonic AuNPs. (a) Schematic illustration of the TA spectroscopy setup with CPL excitation. (b) Jablonski energy-level diagram illustrating the three contributions to the TA signal: excited-state absorption (ESA_1_ and ESA_2_), ground-state bleaching (GSB), and stimulated emission (SE). (c) TA spectra of the AuNPs under LCP excitation at representative time delays. (d, e) TA profiles at 490 nm (ESA_2_) under LCP/RCP excitation, together with their multi-exponential fitting results.

**Extended data figure 4.**
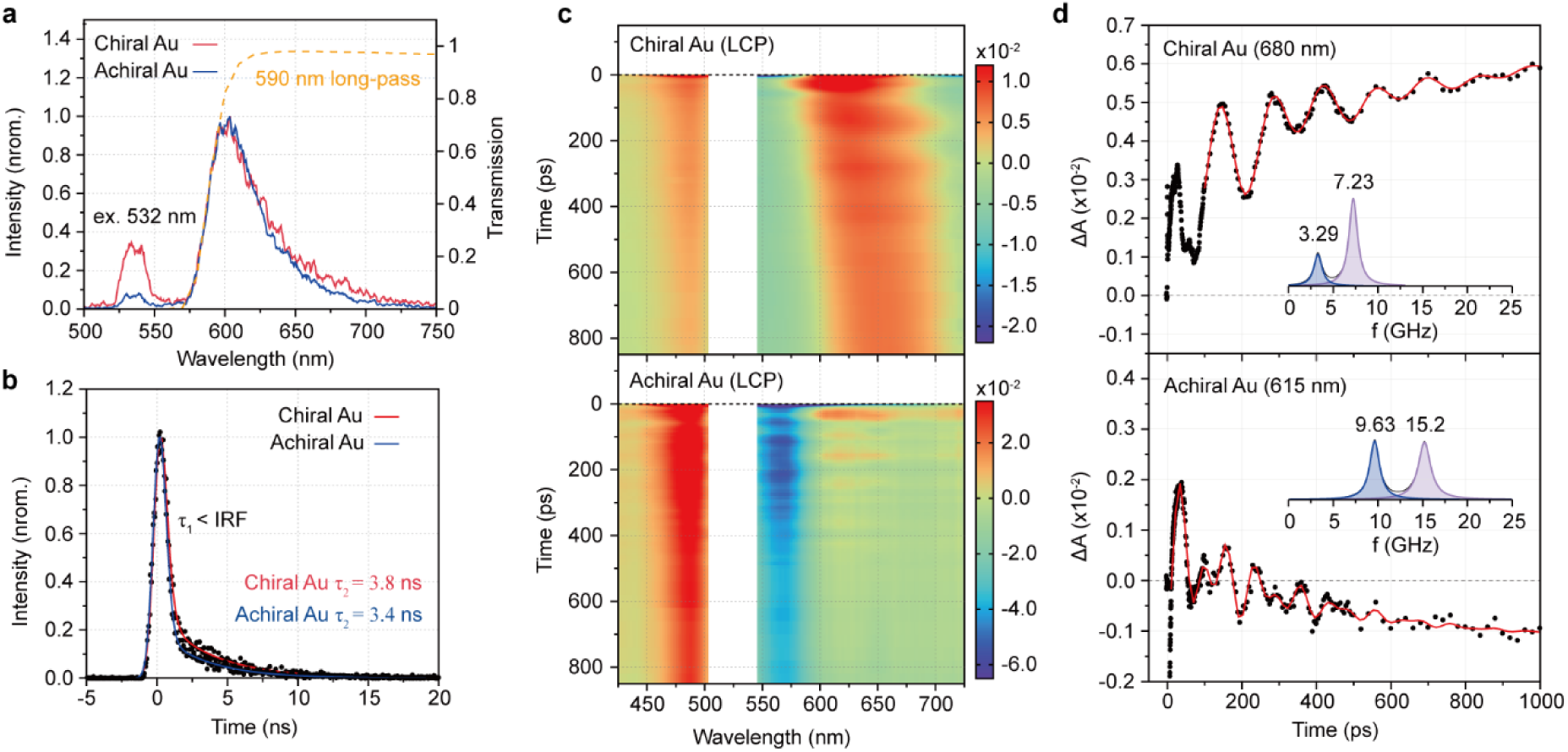
Femtosecond transient absorption (TA) characterization of chiral and achiral plasmonic AuNPs. (a) Time-resolved photoluminescence (TRPL) decay profiles of the AuNPs recorded at 600 nm and (b) corresponding time-gated PL spectra at 5 ns. A 590 nm long-pass filter (dashed orange line) was used to suppress the pump and solvent Raman scatterings. The first decay component (τ_1_) was limited by the width of the instrument response function (IRF). (c) Intensity-scaled TA contour maps of chiral and achiral AuNPs within 1 ns, highlighting apparent coherent phonon oscillations in the TA signals. (d) Linear-prediction singular value decomposition (LPSVD) analysis of the damped oscillations in the TA profiles resolved at the band edges. Two phonon modes were resolved for both samples: the high frequency component is attributed to the breathing motion of the whole nanostructure, whereas the redundant lower frequency component originates from the tip-to-tip acoustic motion of the cubic structure^53^.

**Extended data figure 5.**
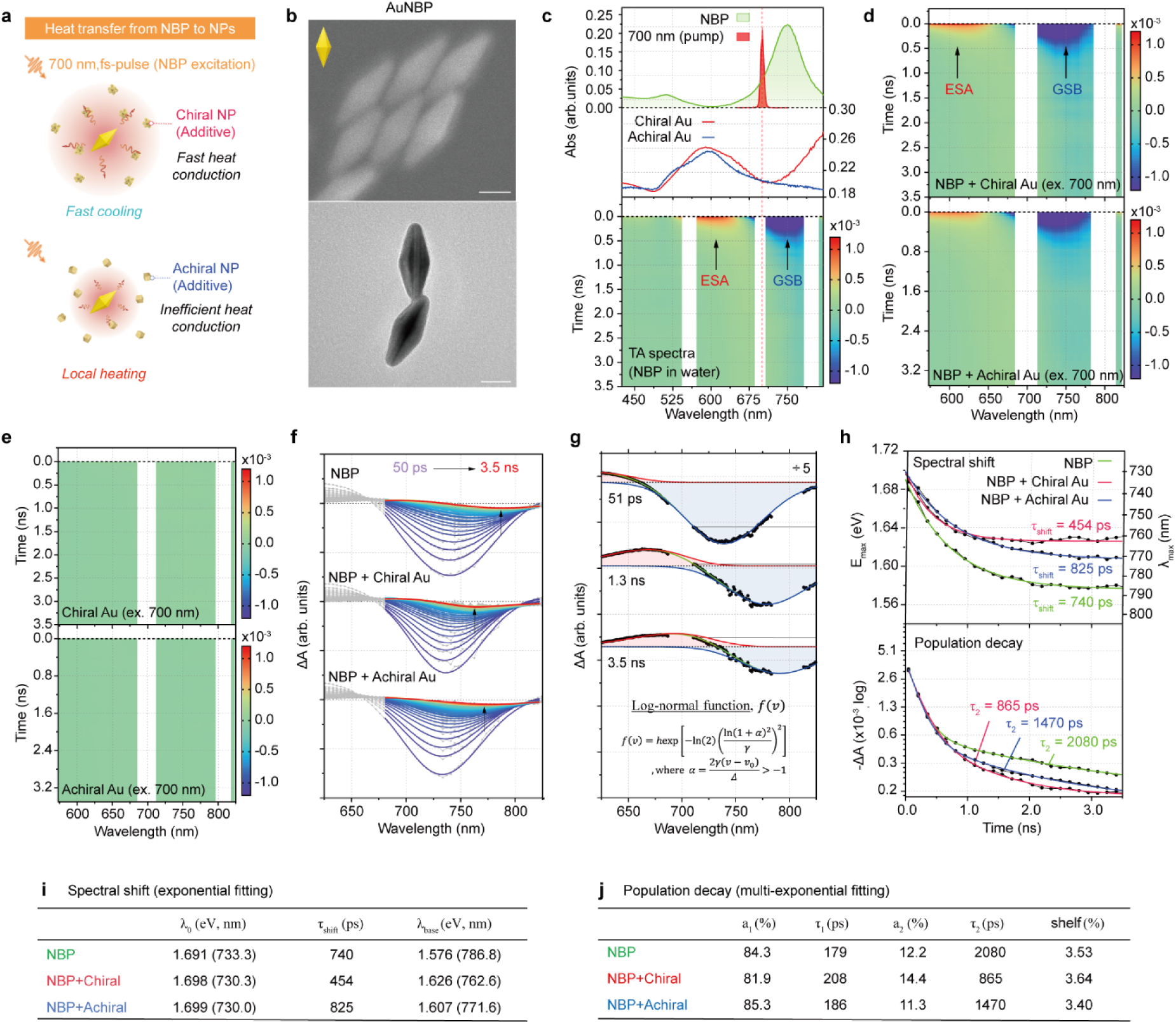
Transient absorption (TA) characterization of Au nano-bipyramids (NBPs) with chiral and achiral AuNP additives. (a) Schematic illustration of the accelerated heat-dissipation process of Au nano-bipyramids (NBPs) in the presence of chiral and achiral AuNP additives. Here, the NBP serves as a spectroscopic probe while the two AuNPs act as heat conductors. (b) SEM (upper) and TEM (lower) images of Au NBPs. (Scale bar: 50 nm). (c) Steady-state absorption spectra of the NBP and chiral/achiral AuNPs in water (top and middle), and TA contour map of the NBP under 700 nm excitation (bottom). The 700 nm excitation profile (red-shaded curve) is overlaid. In the TA map, strong scattering from the pump (700 nm), fundamentals (800 nm) were removed. (d) TA contour maps of the NBP solutions containing chiral and achiral AuNPs, showing a strong negative GSB signal around 750 nm. (e) TA contour maps of chiral and achiral AuNPs under 700 nm excitation, showing negligible TA signal over the entire spectral range. (f) Raw TA spectra of the NBP and the two mixtures containing the AuNP additives at representative time-delays (150 ps intervals). For clarity, global fitting results using log-normal functions are overlaid. Vertical arrows indicate the center positions of the deconvoluted GSB band at 3.5 ns. (g) Global fitting analysis of the truncated TA spectra of the NBP. Three representative TA spectra are shown together with the global fitting curves. Two log-normal functions were used to deconvolute the GSB (blue) contribution from the ESA (red), where *h*, *ν*_0_, *Δ*, and *γ* denote peak height, peak center (in eV), width, and asymmetry parameter, respectively. During the global fitting procedure, the width and asymmetry parameters were shared across the entire data set. (h) Temporal changes in the center position (*ν*_0_: E_max_ in eV, or λ_max_ in nm) and amplitude (*h*) of the deconvoluted GSB band, analyzed using multi-exponential decay functions. (i, j) Exponential fitting results for the dynamic spectral shift, *ν*_0_(*t*), and population decay, *h*(*t*), of the deconvoluted GSB band.

**Extended data figure 6.**
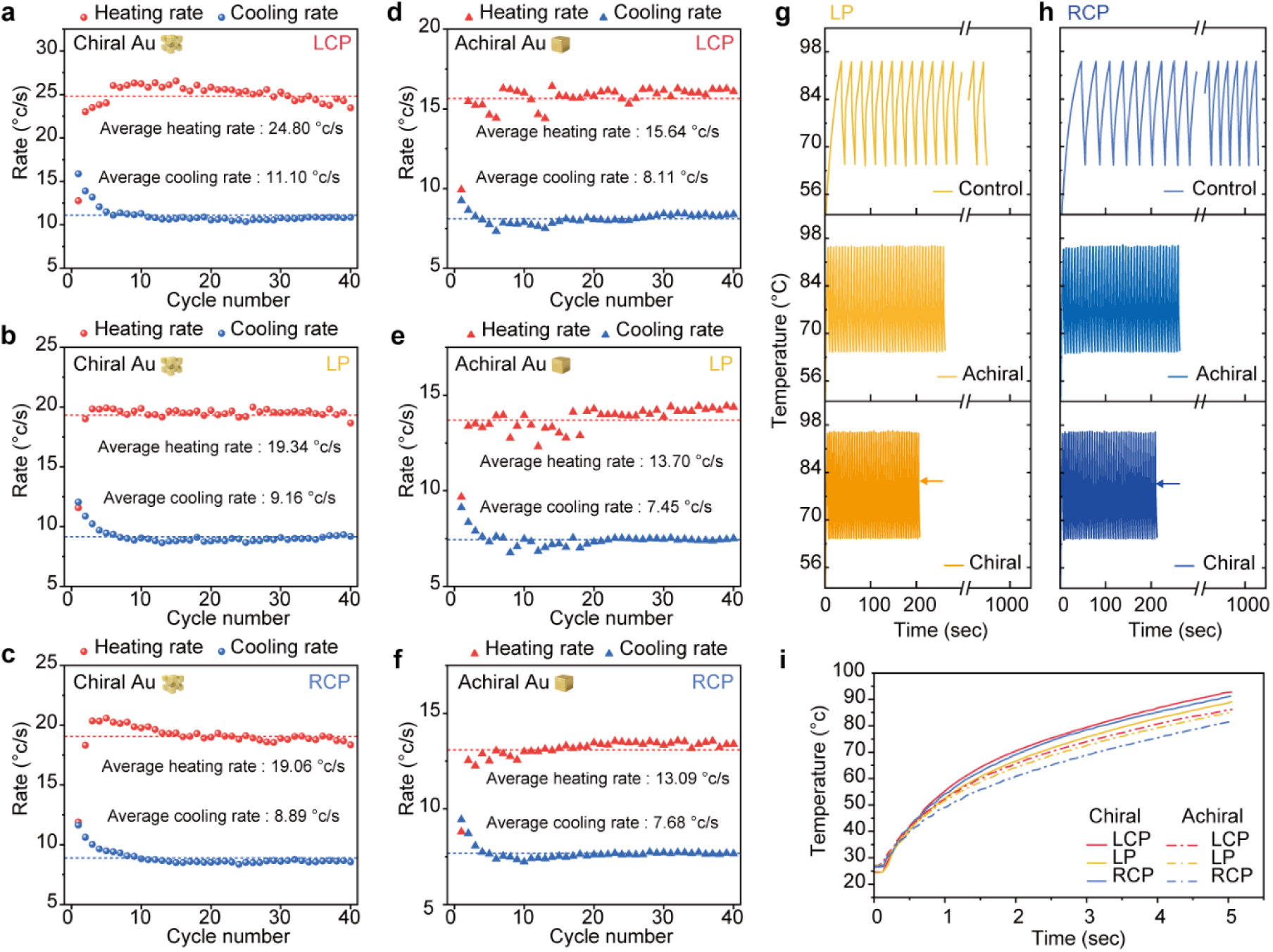
Photothermal cycling performance of chiral and achiral AuNPs under CPL (532 nm). (a–c) Cycle-by-cycle heating and cooling rates of chiral AuNPs during 40 photothermal cycles under (a) LCP, (b) LP, and (c) RCP irradiation, respectively. (d–f) Cycle-by-cycle heating and cooling rates of achiral AuNPs under (d) LCP, (e) LP and (f) RCP irradiation, showing lower average heating and cooling rates than the chiral AuNPs under the corresponding polarization conditions. (g,h) Real-time photothermal cycling profiles of deionized water (control), achiral AuNPs and chiral AuNPs under (g) LP and (h) RCP irradiation, showing that chiral AuNPs enable faster thermal cycling than achiral AuNPs under both polarization conditions. (i) Initial photothermal heating profiles of chiral and achiral AuNPs under LCP, LP, and RCP irradiation, used to evaluate the light-to-heat conversion behavior and calculate photothermal conversion efficiency.

**Extended data figure 7.**
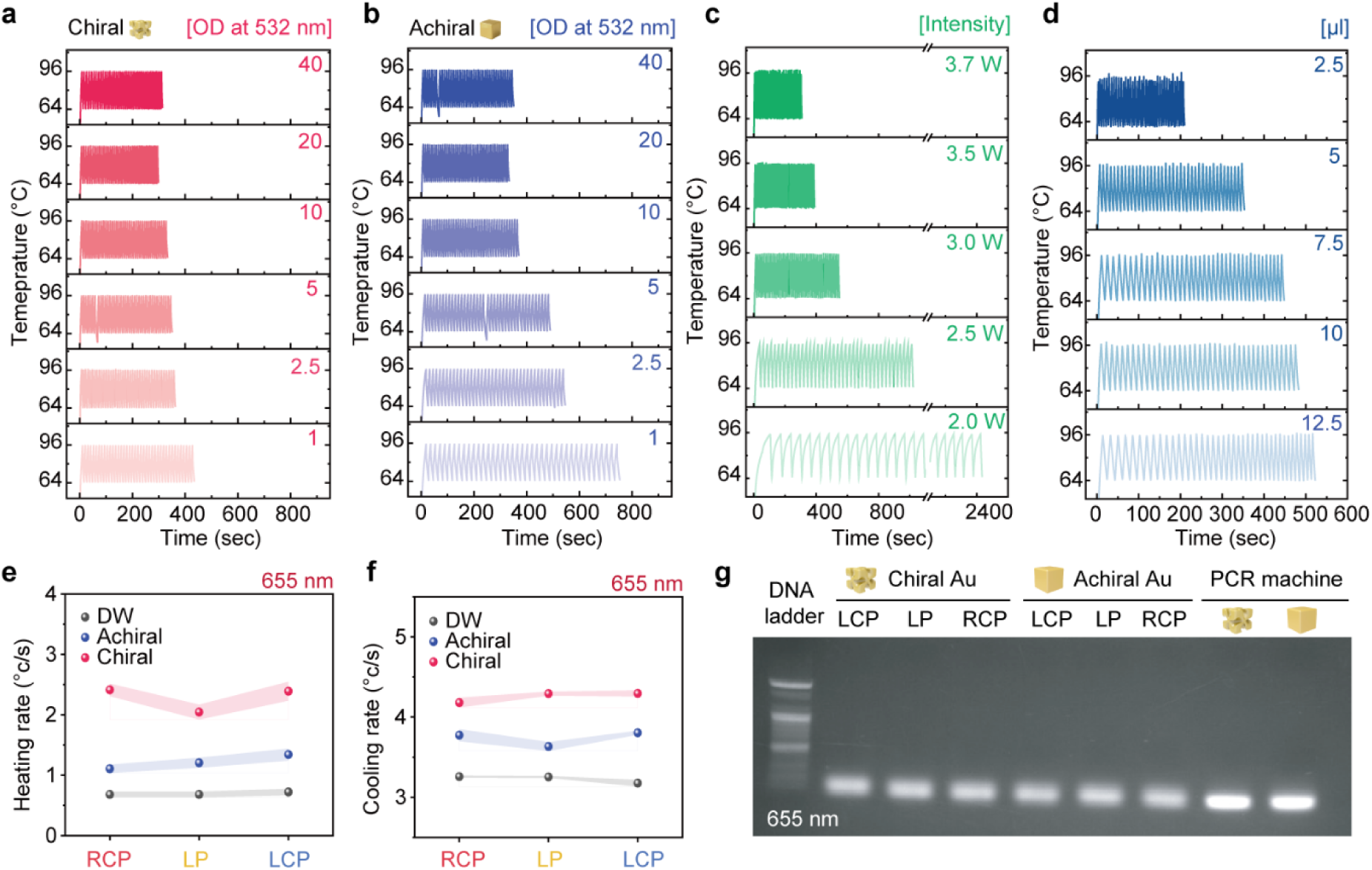
Optimization of photothermal temperature cycling conditions for Au nanoparticle-mediated photonic PCR. (a, b) Temperature profiles over 40 heating–cooling cycles obtained with different optical densities of (a) chiral and (b) achiral AuNPs at 532 nm. The optical density varied from OD 1 to 40 to evaluate the concentration-dependent photothermal response. (c) Temperature cycling profiles of chiral AuNPs under different 532 nm laser powers ranging from 2.0 to 3.7 W. (d) Temperature cycling profiles obtained with different volumes of chiral Au nanoparticle solution, from 2.5 to 12.5 μl, while maintaining a total reaction volume of 10 μl. (e, f) Comparison of (e) heating and (f) cooling rates of deionized water, achiral AuNPs, and chiral AuNPs under RCP, LP, and LCP irradiation at 655 nm. Shaded regions indicate the standard deviation from *n* = 3 independent measurements. (g) Gel electrophoresis analysis of PCR products amplified using chiral and achiral Au nanoparticle-assisted photothermal cycling under different polarization conditions at 655 nm, with amplification using a conventional PCR machine included as a control.

**Extended data figure 8.**
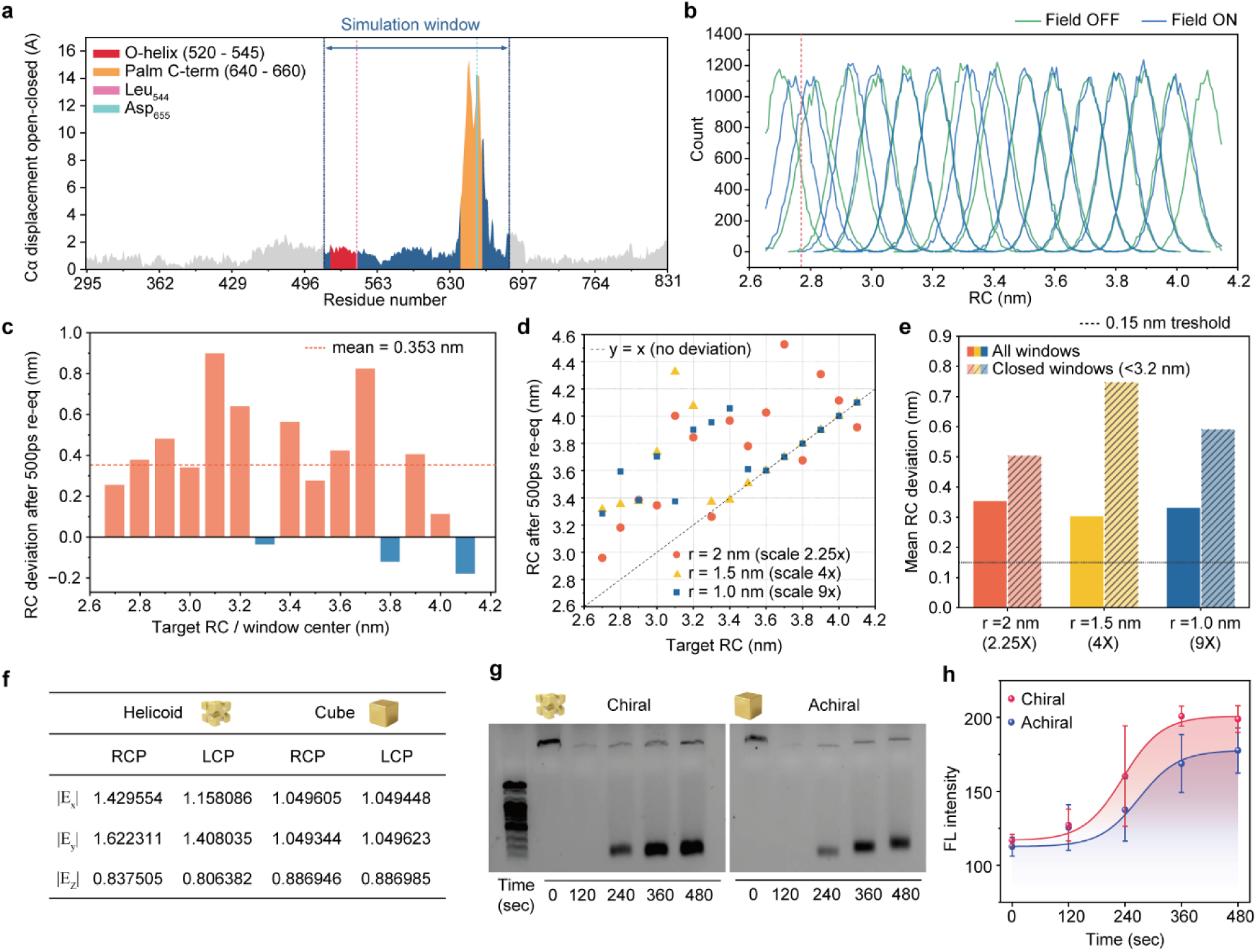
Construction and validation of chiral electromagnetic field-assisted molecular dynamics simulations of Taq DNA polymerase. (a) Residue-level Cα displacement analysis of Taq DNA polymerase during the transition from the open to closed conformational state. The simulation window used for subsequent molecular dynamics analysis is highlighted, together with the O-helix region, palm C-terminal region and the Leu544 and Asp655 residues used to define the reaction coordinate. (b) Histogram overlap analysis of umbrella sampling windows along the Leu544–Asp655 reaction coordinate under field-off and field-on conditions. The continuous overlap between adjacent windows indicates sufficient sampling quality for potential of mean force analysis. (c) Reaction-coordinate deviation after 500 ps reequilibration as a function of target umbrella-window centre. The dashed line indicates the mean deviation across all windows. (d) Reaction coordinate after 500 ps reequilibration plotted against the target reaction coordinate under different field-strength conditions, corresponding to field scaling factors estimated at distances of r = 2.0, 1.5 and 1.0 nm from the nanoparticle surface. (e) Mean reaction-coordinate deviation after 500 ps re-equilibration for all sampling windows and for closed-state windows with reaction coordinates below 3.2 nm. The larger deviation in the closed-state windows under stronger field conditions indicates that the chiral electromagnetic field preferentially destabilizes the closed O-helix conformation and shifts the polymerase toward a more open state. (f) Calculated local electric-field intensities of chiral helicoid and achiral cubic nanostructures. Calculated local electric-field intensity components, |*E_x_*|, |*E_y_*| and |*E_z_*|. Simulations were performed under RCP and LCP plane-wave excitation propagating along the z direction at 532 nm with an incident electric-field amplitude of |*E_inc_*| = 1 V m^−1^. (g) Agarose gel electrophoresis analysis of photonic PCR products amplified on chiral and achiral AuNPs under RCP irradiation at different reaction times. (h) Time-dependent fluorescence intensity profiles of PCR amplification on chiral and achiral AuNPs under RCP irradiation. Error bars represent s.d. from n = 3 independent measurements for each condition.

**Extended data figure 9.**
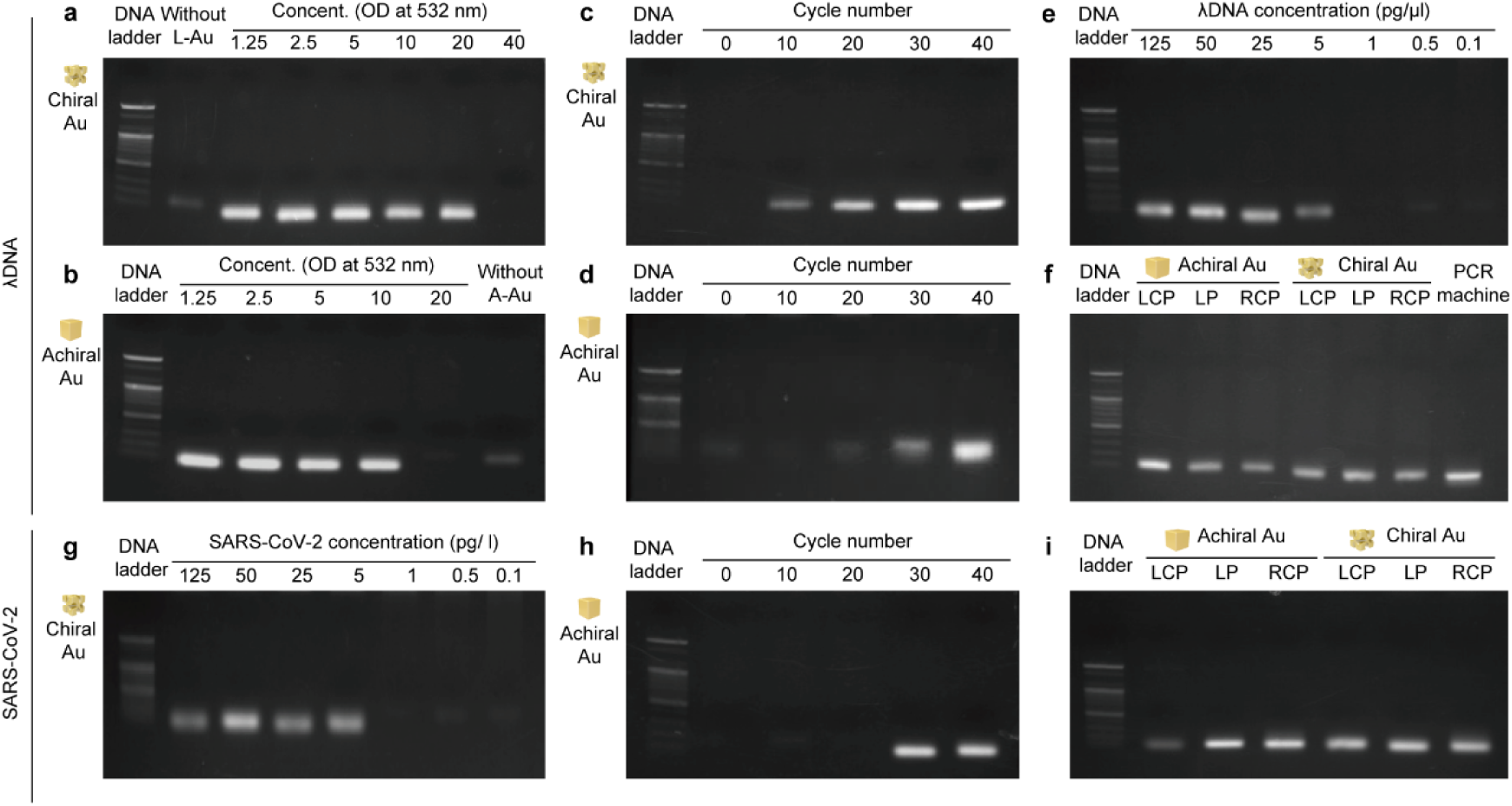
Gel analysis of Photonic PCR products under different amplification conditions. (**a,b**) Amplification of λDNA using chiral (a) and achiral (b) AuNPs at different optical densities (OD at 532 nm). OD values were 1.25, 2.5, 5, 10, 20 and 40 for chiral AuNPs and 1.25, 2.5, 5, 10 and 20 for achiral AuNPs. Lanes labelled “Without L-Au” and “Without A-Au” indicate reactions without the corresponding AuNPs. (**c,d**) Cycle-dependent amplification of λDNA using chiral (c) and achiral (d) AuNPs after 0, 10, 20, 30 and 40 thermal cycles. (**e**) Amplification from serially diluted λDNA templates (125, 50, 25, 5, 1, 0.5 and 0.1 pg μL^−1^) using chiral AuNPs. (**f**) Agarose gel electrophoresis confirming successful λDNA amplification after 30 photonic PCR cycles using achiral and chiral AuNPs under LCP, LP and RCP irradiation. Amplification using a conventional thermal cycler is included as a positive control in the final lane. (**g**) Amplification of the SARS-CoV-2 E-gene target from serially diluted plasmid templates (125, 50, 25, 5, 1, 0.5 and 0.1 pg μL^−1^) using chiral AuNPs. (**h**) Cycle-dependent amplification of the SARS-CoV-2 E-gene target using achiral AuNPs after 0, 10, 20, 30 and 40 thermal cycles. (**i**) Agarose gel electrophoresis confirming successful SARS-CoV-2 E-gene amplification after 30 photonic PCR cycles using achiral and chiral AuNPs under LCP, LP and RCP irradiation.

**Extended data figure 10.**
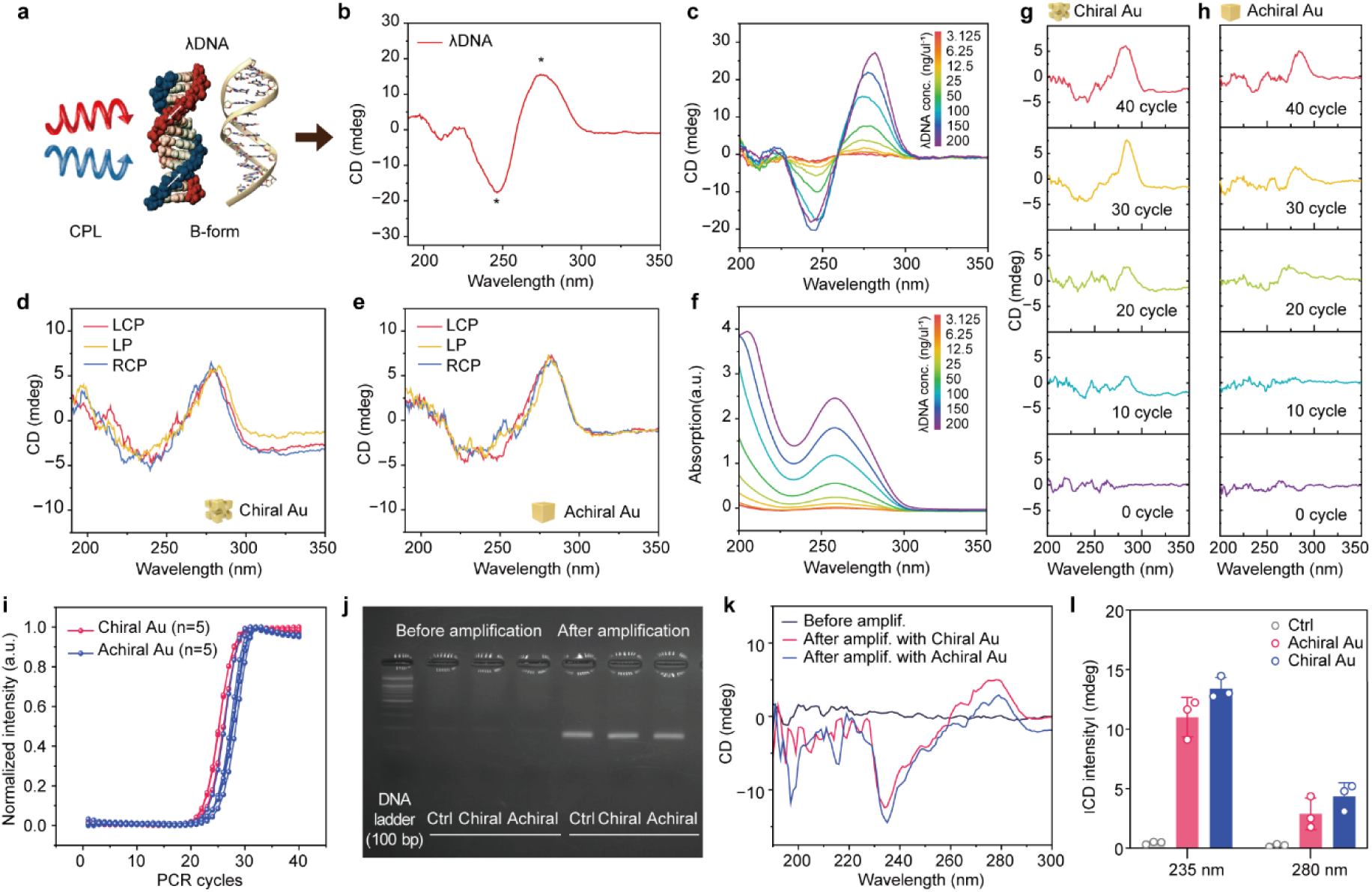
CD analysis of λDNA amplification during photonic PCR on chiral and achiral AuNPs. (a) Schematic illustration of the CD-based analysis of λDNA possessing a double-helical structure. (b) Representative CD spectrum of λDNA, showing the characteristic DNA features with a negative band near 245 nm and a positive band near 275 nm. (c) Concentration-dependent CD spectra of λDNA from 3.125 to 200 ng μL^−1^, demonstrating the gradual increase in CD intensity with increasing DNA concentration. (d,e) CD spectra of PCR products obtained from chiral Au and achiral AuNPs after 30 cycle of PCR under LCP, LP, and RCP irradiation, respectively. (f) Corresponding UV–Vis absorbance spectra reveal increased absorption at ∼260 nm with increasing DNA concentration. (g,h) Cycle-dependent CD spectra of PCR products collected after 0, 10, 20, 30, and 40 thermal cycles on chiral and achiral AuNPs, respectively. The progressive emergence of the CD signature indicates successful DNA amplification, with the chiral Au nanoparticle showing stronger cycle-dependent signal development. (i) Quantitative PCR amplification curves of the β-actin gene using PCR machines embedded with either chiral or achiral AuNPs (OD = 20 at 532 nm). Each condition was tested in quintuplicate (n = 5). (j) Gel electrophoresis image comparing DNA band intensity before amplification and after PCR using machines embedded with either L-AuNP or A-AuNP. (k) CD spectra of DNA samples before and after PCR amplification, demonstrating enhanced chiroptical signals after amplification. (l) CD intensities at 235 nm and 280 nm extracted from CD spectra. Negative CD values at 235 nm are plotted as absolute values.

