## Supplementary Information for "Chirality-Driven Photonic PCR Enables Label-Free Genetic Amplification"

**Supplementary Table 1 | Temperature parameters used to calculate photothermal conversion efficiency.**

| OD = 20 at 532 nm | | $T_{min}$ (°C) | $T_{max}$ (°C) | ΔT (°C) | Time (sec) |
|---|---|---|---|---|---|
| Chiral AuNP | LCP | 24.603 | 92.792 | 68.188 | 5.04 |
| | LP | 24.286 | 89.039 | 64.752 | 5.043 |
| | RCP | 26.485 | 91.207 | 64.72 | 5.027 |
| Achiral AuNP | LCP | 27.097 | 86.181 | 59.084 | 5.047 |
| | LP | 27.030 | 85.011 | 57.981 | 5.022 |
| | RCP | 26.854 | 81.730 | 54.875 | 5.051 |

Average minimum temperature ($T_{min}$), maximum temperature ($T_{max}$), temperature increase ($\Delta T$) and response time measured for chiral and achiral AuNPs under LCP, LP and RCP irradiation. All measurements were performed at OD = 20 at 532 nm. The response time corresponds to the time interval used to determine the initial temperature rise for photothermal conversion efficiency calculation.

**Supplementary Table 2 | Comparison of this work with the previously reported photonic PCR works for heating rates as a function of 30-cycle amplification time. This data are depicted graphically in Fig 3j.**

| | Platform / Material | Light Source | Heating Rate (°C/s) | Cooling Rate (°C/s) | Reaction Volume | Time for 30 Cycles (s) | Reference |
|---|---|---|---|---|---|---|---|
| [1] | Au thin-film photonic PCR | Blue LEDs (450 nm) | 12.79 | 6.60 | 10 µL | 300.0 | *Light Sci. Appl.* **2015**, *4*, e280 |
| [2] | Au nanofilm photonic PCR | Blue LED (447.5 nm) | 4.44 | 2.65 | 10 µL | 600.0 | *Biosens. Bioelectron. X* **2023**, *14* ,100346 |
| [3] | Metaphotonic absorber PCR chip (TiN) | Visible LED | 14.00 | 8.00 | 10 µL | 225.0 | *Adv. Mater.* **2024**, *36*, 2311931 |
| [4] | Plasmonic optical well | Visible LED | 15.00 | 10.00 | Sub-µL optical wells | 180.0 | *Nano Lett.* **2024**, *24*, 1738–1745 |
| [5] | Nanoplasmonic pillar arrays / 10–50 nm Au | White LED | 11.95 | 7.31 | 10 µL | 210.0 | *ACS Appl. Mater. Inter.* **2020**, *12*, 12533–12540 |
| [6] | Au nanopillar plasmonic arrays (100 nm pillars) | Visible LED | 8.75 | 11.67 | Microfluidic chamber (volume not stated) | 247.5 | *Adv. Sci.* **2025**, *12*, 2411849 |
| [7] | $TiO_2$–PDMS nanocomposite PCR reactor | Visible LED | 18.33 | 3.11 | PCR microreactor (volume not stated) | 600.0 | *J. Nanobiotechnol.* **2025**, *23*, 240 |
| [8] | Au bipyramid nanoheaters | NIR laser (808 nm) | 16.60 | 9.40 | 10 µL | 270.0 | *J. Am. Chem. Soc.* **2017**, *139*, 8054–8057 |
| [9] | rGO photothermal microparticles | NIR laser (808 nm) | 11.60 | 8.30 | 10 µL | 300.0 | *ACS Nano*, **2022**, *16*, 20533-20544 |
| [10] | Au nanoshell photothermal PCR | NIR laser (808 nm) | 6.64 | 3.10 | 20 µL PCR mixture | 600.0 | *Sci. Rep.* **2024**, *14*, 3895 |

| | Platform / Material | Light Source | Heating Rate (°C/s) | Cooling Rate (°C/s) | Reaction Volume | Time for 30 Cycles (s) | Reference |
|---|---|---|---|---|---|---|---|
| [11] | Magneto-optical PCR: $Zn_{0.4}Fe_{2.6}O_4$ core / 12 nm Au shell | NIR laser (808 nm) | 8.00 | 5.00 | 10 µL | 300.0 | *Nat. Biomed. Eng.* **2020**, *4*, 1159–1167 |
| [12] | Spherical AuNP plasmonic thermocycler | NIR laser (808 nm) | 6.10 | 3.10 | 20 µL | 600.0 | *Analyst*, **2012**, *137*, 4475–4481 |
| [13] | Au nanofilm microfluidic chip | Two blue LEDs (top/bottom) | 7.37 | 1.91 | Droplet microreactors (total chamber volume not stated) | 780.0 | *Sci. Rep.* **2021**, *11*, 23338 |
| [14] | Plasmonic thermocycler PoM cartridge | White LED | 18.85 | 8.89 | 2 µL | 300 | *ACS nano*, **2023**, *17*, 6507-6518 |
| [15] | Carbon-black/graphene film | White LED | 12 | 7 | 2 µL | 300 | *Adv. Sci.* **2023**, *10*, 2302072 |
| [16] | Plasmonic cavity membrane enrichment + PCR | Visible LED | 12 | 7 | Several nL in membrane cavities | 300 | *Adv. Mater.* **2024**, *36*, 2403896 |
| [17] | Polymer functionalized Au-film photothermal dPCR | Visible LED | 10 | 10 | Digital microwell array; volume not stated | 400 | *ACS Nano* **2025**, *19*, 27280–27293 |
| [18] | Honeycomb-patterned 160 nm Au-film solid-phase photonic PCR | 448 nm LED, max. 1030 mW | 13.84 | 8.97 | 8 µL | 193.0 | *Adv. Funct. Mater.* **2024**, *34*, 2402972 |
| [19] | Chiral AuNP | CPL laser (532 nm) | 24.80 | 11.10 | 20 µL | 171.7 | *This work* |
| [20] | Achiral AuNP | CPL laser (532 nm) | 15.64 | 8.11 | 20 µL | 238.0 | *This work* |

**Supplementary Table 3 | Oligo nucleotide sequence used for photonic PCR amplification.**

| Target | Specification | Oligonucleotide sequence (5’ to 3’) |
|---|---|---|
| λDNA | Forward primer | GTT CTG AGG TCA TTA CTG GAT C |
| | Backward primer | TAC GCT CCT GTC CGG CAA A |
| SARS CoV-2 | Plasmid for SARS-CoV-2 envelope (E) protein, (113bp) | ACAGGTACGTTAATAGTTAATAGCGTACTTCTTT TTCTTGCTTTCGTGGTATTCTTGCTAGTTACACTA GCCATCCTTACTGCGCTTCGATTGTGTGCGTACT GCTGCAATAT |
| | Forward primer | ACAGGTACGTTAATAGTTAATAGCGT |
| | Backward primer | ATATTGCAGCAGTACGCACACA |

Primer sequences and target nucleotide information used for λDNA and SARS-CoV-2 E gene amplification. The table lists the forward and reverse primer sequences for λDNA, the 113-bp SARS-CoV-2 E gene target sequence inserted in the plasmid template and the corresponding primer sequences used for SARS-CoV-2 target amplification. All sequences are shown in the 5′ to 3′ direction.
